# Epigenetic derepression of FKBP5 by aging and stress contributes to NF-ĸB-driven inflammation and cardiovascular risk

**DOI:** 10.1101/484709

**Authors:** Anthony S. Zannas, Meiwen Jia, Kathrin Hafner, Jens Baumert, Tobias Wiechmann, Julius C. Pape, Janine Arloth, Maik Ködel, Silvia Martinelli, Maria Roitman, Simone Röh, Andreas Haehle, Rebecca T. Emeny, Stella Iurato, Tania Carrillo-Roa, Jari Lahti, Katri Räikkönen, Johan G. Eriksson, Amanda J. Drake, Melanie Waldenberger, Simone Wahl, Sonja Kunze, Susanne Lucae, Bekh Bradley, Christian Gieger, Felix Hausch, Alicia K. Smith, Kerry J. Ressler, Bertram Müller-Myhsok, Karl-Heinz Ladwig, Theo Rein, Nils C. Gassen, Elisabeth B. Binder

## Abstract

Aging and psychosocial stress are associated with increased inflammation and disease risk, but the underlying molecular mechanisms are poorly understood. Because both aging and stress are also associated with lasting epigenetic changes, a plausible hypothesis is that stress exposure along the lifespan could confer disease risk by epigenetically deregulating molecules involved in inflammatory processes. Here, by combining large-scale analyses in human cohorts with mechanistic *in vitro* investigations, we found that FKBP5, a protein implicated in stress physiology, contributes to these relations. Across independent human cohorts (total n=3,131), aging and stress-related phenotypes were synergistically associated with epigenetic derepression of *FKBP5*. These age/stress-related epigenetic effects were recapitulated in an *in vitro* model of replicative senescence, whereby we exposed replicating human fibroblasts to stress (glucocorticoid) hormones. Unbiased genome-wide analyses in human blood linked higher *FKBP5* mRNA with a proinflammatory profile and altered NF-κB-related gene networks. Accordingly, experiments in immune cells showed that *FKBP5* overexpression promotes inflammation by strengthening the interactions of NF-κB regulatory kinases, whereas opposing FKBP5 either by genetic deletion (CRISPR/Cas9-mediated) or selective pharmacological inhibition prevented the effects on NF-κB. Further, the age/stress-related epigenetic signature enhanced *FKBP5* responsivity to NF-κB through a positive feedback loop and was present in individuals with a history of acute myocardial infarction, a disease state linked to peripheral inflammation. These findings suggest that FKBP5-NF-κB signaling mediates inflammation associated with aging and stress, potentially contributing to cardiovascular risk, and may thus point to novel biomarker and treatment possibilities.

**Significance:** Diseases of the aging are the leading cause of morbidity and mortality. Elucidating the molecular mechanisms through which modifiable factors, such as psychosocial stress, confer risk for aging-related disease can have profound implications. Here, by combining studies in humans with experiments in cells, we find that aging and stress synergize to epigenetically derepress FKBP5, a protein implicated in stress physiology. Higher FKBP5 promotes inflammation by activating the master immune regulator NF-κB, whereas opposing FKBP5 – either genetically or pharmacologically– prevents the effects on NF-κB. Further, the age/stress-related epigenetic signature of *FKBP5* is associated with history of myocardial infarction, a disease state linked to inflammation. These findings provide molecular insights into stress-related disease and may point to novel biomarker and treatment possibilities.

## Introduction

Aging is the single most important risk factor for several disease phenotypes that are leading causes of morbidity and mortality (1). Yet individuals of the same age exhibit substantial variability in their propensity to develop aging-related disease (2). Among important factors influencing disease risk, studies show that psychosocial stressors, such as childhood trauma, as well as stress-related psychiatric disorders, including major depressive disorder (MDD), increase risk for aging-related diseases, most notably cardiovascular syndromes (3-7). Studies further suggest that aging and stress-related phenotypes may together confer disease risk by increasing peripheral inflammation (5, 8-11), but the underlying mechanisms and cascade of molecular events are poorly understood.

Mechanistically, the effects of stress on inflammation and disease risk could be driven by stress-responsive molecules able to modulate immune function. A plausible such molecule to examine is the FK506-binding protein 51 (FKBP51/FKBP5), a protein co-chaperone that is acutely induced by stress and can influence downstream biological processes through protein-protein interactions (12-19). Interestingly, FKBP5 upregulation has been observed not only upon stress exposure and glucocorticoid stimulation, but also in the aging brain (20, 21) and in some disease phenotypes (15, 17, 20, 22). Yet it is unknown whether aging regulates FKBP5 in the immune system and how this effect, if present, could shape risk for cardiovascular disease. Both aging and stress can have lasting effects on the epigenome (23-26), and *FKBP5* transcription can be regulated by epigenetic mechanisms (27-29); thus, a plausible hypothesis is that stress exposure along the lifespan could epigenetically deregulate *FKBP5* in immune cells, potentially contributing to peripheral inflammation and disease risk.

Here we address these questions by combining genome-wide analyses in human cohorts with mechanistic *in vitro* investigations. Convergent findings support a model whereby aging and stress-related phenotypes synergize to decrease DNA methylation at selected enhancer-related *FKBP5* sites, epigenetically derepressing *FKBP5* in whole blood and in distinct immune cell subtypes. Derepressed *FKBP5* in turn promotes NF-κB (nuclear factor kappa-light-chain-enhancer of activated B cells)-driven peripheral inflammation. Accordingly, the age/stress-related *FKBP5* epigenetic signature is present in individuals with a history of acute myocardial infarction, a disease state linked to peripheral inflammation. We further find that the cellular effects of stress on NF-κB are prevented by either CRISPR/Cas9 deletion of the *FKBP5* gene or a selective FKBP5 antagonist, suggesting that FKBP5-NF-κB signaling is a tractable treatment candidate. Together these findings provide molecular insights into mechanisms linking aging and stress-related phenotypes with peripheral inflammation and cardiovascular risk, thereby pointing to novel biomarker and intervention possibilities.

## Results

### FKBP5 DNA methylation decreases along the lifespan at selected CpGs

DNA methylation at cytosine-guanine dinucleotides (CpG) has been shown to change with age (25), possibly moderated by environmental factors, including psychosocial stress (24, 30). These age-related epigenetic changes are thought to contribute to interindividual variability in genomic function and disease risk (31, 32). To identify *FKBP5* CpG sites that may be subject to this phenomenon, we first examined how *FKBP5* DNA methylation changes throughout life, using Illumina HumanMethylation450 BeadChip (450K) data from three independent cohorts with broad age range and documented stress-related phenotypes: the Grady Trauma Project (GTP; n = 393, age range 18-77 years); the Cooperative Health Research in the Region of Augsburg F4 community study (KORA; n = 1,727, age range 32-81 years); and the Max Planck Institute of Psychiatry cohort (MPIP; n = 537, age range 18-87 years) (demographics in Supplementary Table 1). These analyses included all available CpGs covered by the 450K within or in close proximity (10kb upstream or downstream) to the *FKBP5* locus (chromosome 6p21.31). After controlling for potential confounders including blood cell type heterogeneity (see Methods) and after FDR correction for multiple comparisons, two CpGs (cg20813374 and cg00130530) showed consistent and robust age-related decrease in methylation across all cohorts (detailed statistics in Supplementary Table 2). These two age-related sites lie in close proximity to each other proximally (< 500 bp) upstream of the *FKBP5* transcription start site (TSS; -462bp for cg20813374 and -484bp for cg00130530; UCSC Genome Browser; Supplementary Table 2) and show significant pairwise correlations in all cohorts (GTP: r = 0.83, p < 2.2 x 10^-16^; KORA: r = 0.61, p < 2.2 x 10^-16^; MPIP r = 0.37, p < 2.2 x 10^-16^). The association of age with average methylation of the two CpGs is depicted in Fig. 1A and Supplementary Fig. 1A. To validate this finding with a non-hybridization-based DNA methylation method, we performed targeted bisulfite sequencing with the Illumina MiSeq in a smaller sample of female subjects, again observing robust pairwise correlation of the two CpGs (r = 0.62, p = 5.7 x 10^-10^) and significantly lower average methylation of the two sites with increasing age (n = 77, p = 1.9 x 10^-2^; Supplementary Fig. 2). Given the close proximity and the consistent pairwise correlations between the two age-related *FKBP5* CpGs, all subsequent analyses examined the average methylation levels of the two sites.

**Figure 1.**
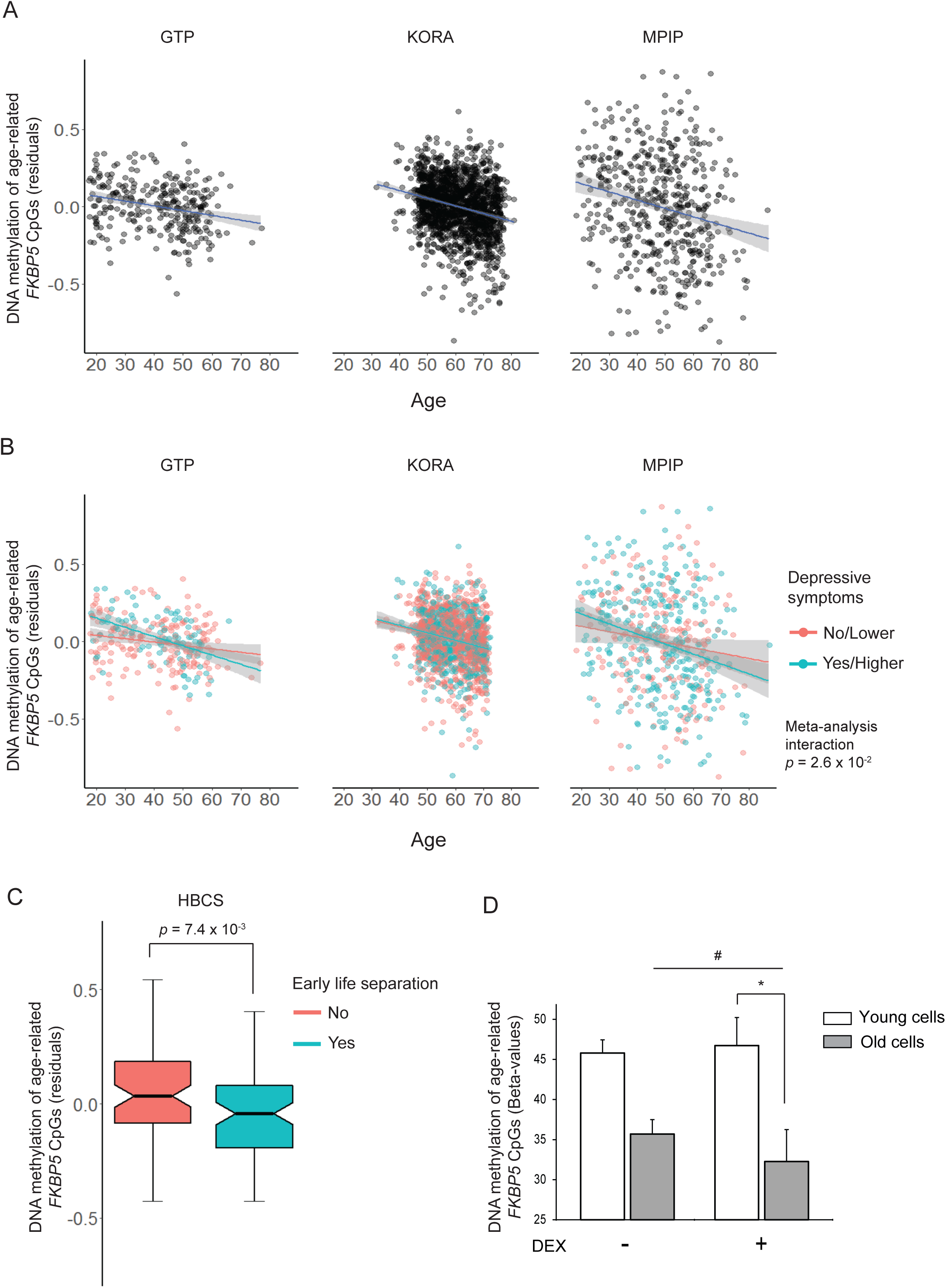
Aging and stress are together associated with decreased DNA methylation at selected *FKBP5* CpGs. **(A)** Methylation decreases at selected *FKBP5* CpGs along the human lifespan (GTP: β_age_ = -0.0045, SE = 0.0008, p = 8 x 10^-8^; KORA: β_age_ = -0.0055, SE = 0.0005, p < 2 x 10^-16^; MPIP: β_age_ = -0.0064, SE = 0.0012, p = 7 x 10^-8^; total n = 2,523). **(B)** Depressive phenotypes are associated with accelerated age-related *FKBP5* demethylation (total n = 2,249, meta-analysis interaction p = 2.6 x 10^-2^, heterogeneity p = 2.7 x 10^-1^). Statistics per cohort: GTP: interaction p = 1.9 x 10^-2^, β_age_ for moderate/severe depression = -0.0075 (SE = 0.0014) *vs.* β_age_ for no/mild depression = -0.0032 (SE = 0.0011); KORA: interaction p = 6.3 x 10^-1^, β_age_ for higher levels of depression = -0.0063 (SE = 0.0011) *vs.* β_age_ for lower levels of depression = -0.0047 (SE = 0.0007); MPIP: interaction p = 1.9 x 10^-1^, β_age_ for depressed = -0.0077 (SE = 0.0015) *vs.* β_age_ for non-depressed = -0.0044 (SE = 0.0019). **(C)** Early life separation is associated with demethylation of the age-related *FKBP5* CpGs in the HBCS (βseparation = - 0.0932, SE = 0.0343, p = 7.4 x 10^-3^, mean DNA methylation difference = 1.4%). All coefficients and p values are derived from linear regression models using M-values for DNA methylation and after correcting for potential confounders (see Methods). **(D)** *In vitro* aging and exposure to the stress hormone (glucocorticoid) receptor agonist dexamethasone (DEX) additively decrease methylation at the age-related *FKBP5* CpGs in the IMR-90 fibroblast model of replicative senescence (F_1,6_ = 6.3, interaction p = 4.6 x 10^-2^, n = 4 replicates per age group). Statistical comparisons were performed with two-way mixed-design ANOVA (according to experimental design), using replicative age as the between-subject and DEX treatment as the within-subject factor. Statistically significant effects were followed with Bonferroni-corrected pairwise comparisons, shown as follows: * p < 5 x 10^-2^, statistically significant pairwise comparisons for young *vs.* old replicative age; # p < 5 x 10^-2^, statistically significant pairwise comparison for vehicle *vs.* DEX-treated old cells. Error bars depict the standard error around the group mean. The y axes in panels (A), (B), and (C) depict the average DNA methylation levels of the two age-related *FKBP5* CpGs (cg20813374 and cg00130530), after adjustment for the respective covariates for each cohort; for a more intuitive visualization, selected panels are also depicted as % DNA methylation (Beta-values in Supplementary Fig. 1). The y axis in panel (D) depicts the average % DNA methylation (Beta-values) of the two *FKBP5* CpGs. GTP, Grady Trauma Project; HBCS, Helsinki Birth Cohort Study; KORA, Cooperative Health Research in the Region of Augsburg F4 community study; MPIP, Max Planck Institute of Psychiatry depression case/control study.

### Age-related decrease in FKBP5 methylation is not confounded by blood cell type heterogeneity and occurs in purified immune cell subtypes

Peripheral blood cell counts change along the lifespan (33), raising the possibility that heterogeneity in blood cell type composition could be confounding the observed inverse relation between age and *FKBP5* methylation levels (34), even when controlling for cell type heterogeneity in the regression model (consistent associations after controlling for calculated blood cell proportions are shown in Supplementary Table 2 and Fig. 1A). To address this possibility, we first performed a series of sensitivity analyses in our cohorts. The inverse relation between age and methylation of the two *FKBP5* CpGs (cg20813374 and cg00130530) was consistent across the GTP, KORA, and MPIP cohorts (all p values < 10^-7^; Fig. 1A), and there was no consistent relation between estimated blood cell subtypes (the potential confounder) and either age or *FKBP5* methylation levels (our variables of interest), suggesting that strong confounding by cell subtypes was not present (Supplementary Table 3). This was further validated using an additional dataset of male and female subjects (n = 213) with both 450K data and differential blood counts; methylation of the age-related CpGs did not significantly correlate with any of the counted blood cell types (Supplementary Table 3) and was again robustly associated with age after adjustment for sex and all cell types (β = -0.0077, SE = 0.0009, p = 5.2 x 10^-15^).

To further rule out potential confounding by cell type heterogeneity and to understand how aging influences *FKBP5* methylation in specific immune cell types, we analyzed publicly available DNA methylation data in whole blood, as well as FACS-sorted CD4 T cells and neutrophils, from male subjects with a broad age range (35). In line with our previous findings, we again observed an inverse relation between age and methylation of the two *FKBP5* CpGs in whole blood (n = 184, r = -0.30, p = 3.6 x 10^-5^). More importantly, the same effect size was present in purified CD4 T cells (n = 46, r = -0.32, p = 3.3 x 10^-2^), whereas this effect was in the same direction but non-significant in purified neutrophils (n = 48, r = -0.20, p = 1.7 x 10^-1^; Supplementary Fig. 3). Together, these findings show that increasing age is associated with lower *FKBP5* methylation in T cell subtypes (and likely other immune cell subtypes) and that this effect is not solely the result of age-related changes in blood cell type composition.

### Childhood trauma and depressive phenotypes are associated with accelerated age-related decrease in FKBP5 methylation

*FKBP5* responds to stress and glucocorticoids and can undergo decrease in DNA methylation at selected stress-responsive CpGs (27-29, 36, 37). Therefore, it is plausible that higher stress burden throughout life could induce lasting epigenetic changes, potentially accelerating decrease in methylation of the two age-related *FKBP5* CpGs. To investigate this hypothesis, we examined the combined effects of age and stress-related phenotypes on average methylation of the two sites. As information on current depressive symptoms was available in all three cohorts, we first investigated this phenotype. After adjusting for blood cell proportions and other covariates (see Methods), depressive phenotypes were associated with a significantly accelerated age-related decrease in *FKBP5* methylation in the GTP, KORA, and MPIP cohorts (total n = 2,249, meta-analysis interaction p = 2.6 x 10^-2^; Fig. 1B). Because early life trauma is among the strongest risk factors for developing MDD (5), we further examined whether the effect of depression on age-related decrease in *FKBP5* methylation is moderated by childhood trauma severity as measured with the childhood trauma questionnaire (CTQ) in the GTP. This stratified analysis yielded a significant age-depression interaction in the higher-CTQ (interaction p = 4.6 x 10^-2^) but not the lower-CTQ group (interaction p = 3.3 x 10^-1^) and no main effect of childhood trauma severity (p = 3.7 x 10^-1^). Lastly, to examine whether exposure to a severe and prolonged early childhood stressor itself is associated with lasting decrease in methylation of the age-related CpGs, we compared elderly individuals that had prolonged early life separation from their parents with sex- and age-matched non-separated controls in a fourth cohort, the Helsinki Birth Cohort Study (n = 160, age range 58-69 years, demographics in Table 1). In this cohort, early life separation was associated with reduced methylation of the age-related CpGs (p = 7.4 x 10^-3^; Fig. 1C and Supplementary Fig. 1B).

Together, these findings suggest that childhood trauma and depressive phenotypes may together accelerate the age-related decrease in *FKBP5* methylation in peripheral blood.

### The effects of both aging and stress on FKBP5 methylation are recapitulated in an in vitro model of replicative senescence

The findings presented above identify two *FKBP5* CpGs (cg20813374 and cg00130530) that show a consistent association of lower methylation levels with stress-related phenotypes and increasing age even within specific cell types; however, these findings are inherently limited by the use of human subjects where experimental manipulation is not feasible. To experimentally validate these associations, we used an *in vitro* model of replicative senescence (IMR-90 fibroblasts) to test whether replicative aging and stress—which is commonly modeled in the dish with the stress (glucocorticoid) hormone receptor agonist dexamethasone (DEX) (27, 38)— influences *FKBP5* methylation at these sites. Population doubling level (PDL) was calculated as previously (39), and *FKBP5* methylation was compared between cells of young (PDL = 22) and old (PDL = 42) replicative age that were treated for 7 days with either vehicle (DMSO) or 100nM DEX. In accordance with our *in vivo* findings, *in vitro* aging and DEX additively decreased DNA methylation at the two *FKBP5* CpGs (interaction p = 4.6 x 10^-2^, DNA methylation decrease in old vs. young cells = 10.1%, additional methylation decrease in old cells treated with DEX vs. vehicle = 3.4%). Together with the aforementioned associations in our human cohorts (Fig. 1A-C), these convergent observations show that aging and stress may influence the two *FKBP5* CpGs across different cohorts, distinct cell types, and contexts.

### Decreased methylation at the age/stress-related FKBP5 CpGs is associated with FKBP5 upregulation in peripheral blood

Changes in DNA methylation can shape gene expression, thereby contributing to cellular function and phenotypic expression (40, 41). The age/stress-related *FKBP5* CpGs identified above lie proximally (< 500 bp) upstream of the TSS for all highly expressed isoforms of *FKBP5* and are intronic only for the minimally expressed variant 2 of the gene (UCSC Genome Browser, UCSC Gens Track; GTEx portal). Integrative analysis of chromatin states using ChromHMM (42) showed that the two CpGs colocalize with signatures that are consistent with either enhancers or flanking active TSS in a large number of cell types (see Supplementary Table 4). In immune cells, the CpGs are commonly mapped to either an enhancer or flanking active TSS (see Supplementary Fig. 4). Furthermore, in most cell types the two sites show intermediate levels of methylation and colocalize with H3K4me1 and H3K27me3 signatures (Roadmap Epigenome Browser; shown for immune cell proxy only in Supplementary Fig. 5). This landscape is most consistent with a poised enhancer (43) that upon transcription factor binding could interact and regulate the downstream *FKBP5* TSS.

To examine whether the methylation status of these CpGs influences gene transcription, we used *FKBP5* mRNA data measured in the GTP cohort with Illumina HumanHT-12 v3 and v4 Expression BeadChip arrays (n = 355). DNA methylation levels of the age/stress-related sites were inversely associated with *FKBP5* mRNA levels (p = 1.6 x 10^-2^; Fig. 2A). We found similar negative correlations in publicly available data from breast tissue samples of control female subjects (n = 84, r = -0.26, p = 1.6 x 10^-2^; Supplementary Fig. 6) (44). Since *FKBP5* transcription is robustly induced by glucocorticoids (29, 45) and given that the CpGs are located in predicted poised enhancers (Supplementary Fig. 5), we then speculated that methylation at the age/stress-related *FKBP5* CpGs could moderate the effect of cortisol on *FKBP5* levels. After confirming a robust positive correlation between cortisol and *FKBP5* mRNA (r = 0.41, p = 1 x 10^-12^), we found that the cortisol-*FKBP5* relationship was significantly stronger in individuals with below-as compared to above-median methylation levels in the GTP (interaction p = 1.4 x 10^-3^; Fig. 2B). In addition, the phenotypes associated with lower methylation levels moderated the relationship between cortisol and *FKBP5* mRNA; specifically, this relationship was significantly stronger in older subjects as defined with a median split of age (interaction p = 2.4 x 10^-5^; Fig. 2C) and in individuals with higher levels of depressive symptoms and childhood trauma exposure (interaction p = 7.3 x 10^-5^; Fig. 2D). These findings are in line with previous observations that stressors can exert lasting epigenetic effects on other sites of the *FKBP5* locus (27-29), and they suggest that the effects of aging and stress-related phenotypes converge at distinct susceptible CpGs to epigenetically derepress *FKBP5* in human peripheral blood.

**Figure 2.**
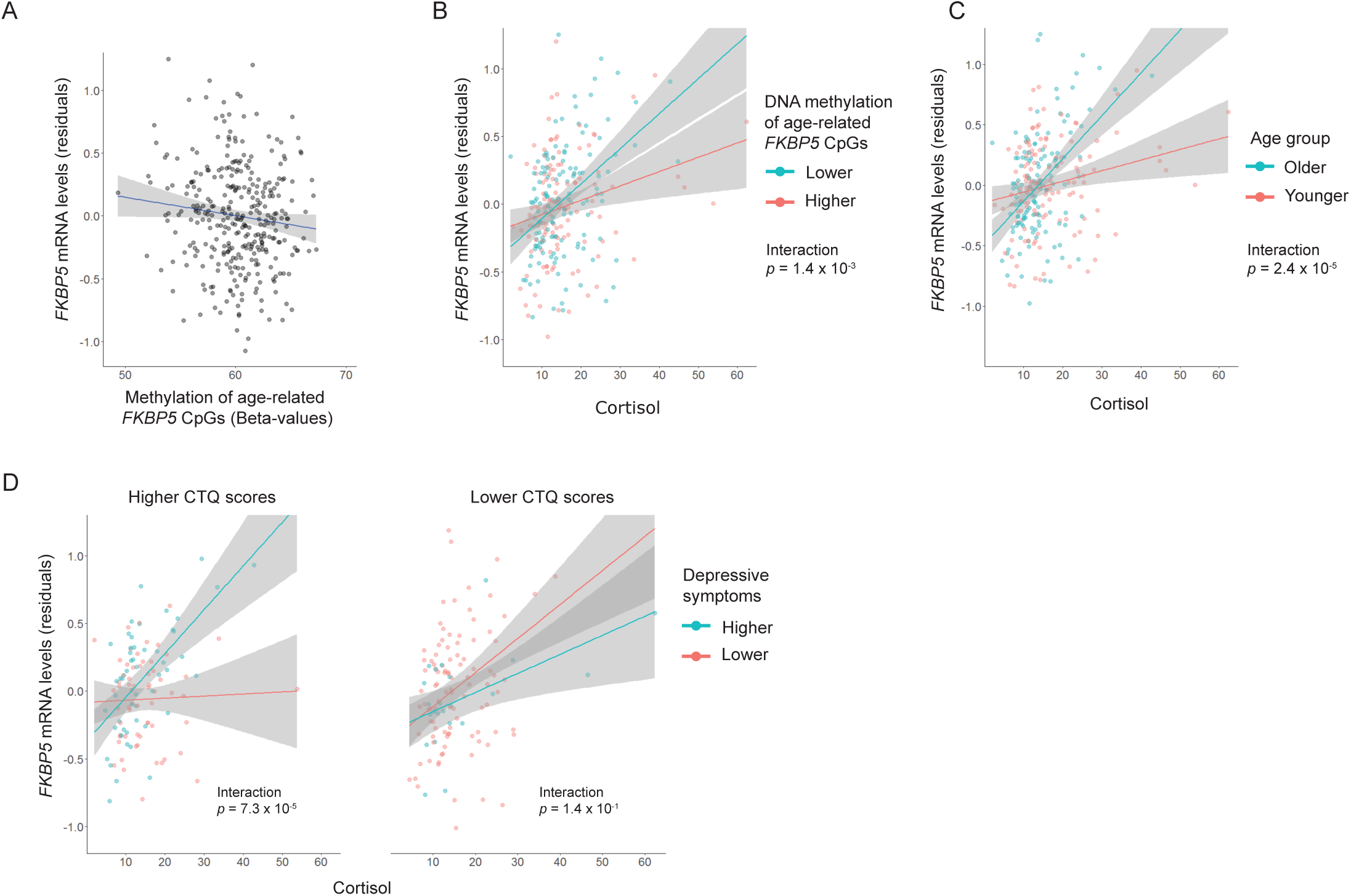
Aging and stress-related phenotypes are associated with epigenetic upregulation of *FKBP5* in peripheral blood in the Grady Trauma Project (n = 355). **(A)** *FKBP5* expression levels are negatively associated with average methylation of the age-related sites (β = -0.3835, SE = 0.1585, p = 1.6 x 10^-2^). **(B** and **C)** The cortisol-*FKBP5* relationship is stronger at lower methylation levels of the age-related *FKBP5* CpGs: interaction p = 1.4 x 10^-3^, β_cortisol_ for lower methylation = 0.0299 (SE = 0.0044) *vs.* β_cortisol_ for higher methylation = 0.0069 (SE = 0.0039). The cortisol-*FKBP5* relationship is stronger in older ages: interaction p = 2.4 x 10^-5^, β_cortisol_ for older subjects = 0.0376 (SE = 0.0050) *vs.* β_cortisol_ for younger subjects = 0.0075 (SE = 0.0035). **(D)** Higher levels of depressive symptoms are associated with stronger cortisol-*FKBP5* relationship in subjects with higher levels of childhood trauma (cortisol-depression interaction p = 7.3 x 10^-5^) but not in subjects with lower levels of childhood trauma (cortisol-depression interaction p = 1.4 x 10^-1^) as defined with the Childhood Trauma Questionnaire (CTQ). Panel (A) depicts the average % methylation levels (Beta-values) of the two age-related *FKBP5* CpGs (cg20813374 and cg00130530).

### FKBP5 upregulation promotes NF-κB-related peripheral inflammation and chemotaxis

To examine the possible functional effects of *FKBP5* deregulation in an unbiased manner, we used genome-wide gene expression data from peripheral blood in the GTP cohort (n = 355) to identify genes that are co-regulated with *FKBP5*. After FDR correction for multiple comparisons (FDR-adjusted p < 0.05), *FKBP5* correlated significantly with a total of 3,275 genes (Supplementary Table 5). Using these transcripts as input and the unique array genes expressed above background (except *FKBP5*) as reference (9,538 genes), we performed pathway and disease association analysis in WebGestalt. The strongest enrichment was observed for inflammation and was conferred by a total of 123 inflammation-related genes (FDR-adjusted p = 8.1 x 10^-6^; Fig. 3A; Supplementary Table 6). Notably, *FKBP5* showed robust positive correlations with a host of proinflammatory genes, such as interleukin and toll-like receptors (Supplementary Table 7). Furthermore, *FKBP5* levels were positively associated with the granulocyte proportion (r = 0.21, p = 8.4 x 10^-5^) and the granulocyte to lymphocyte (G/L) ratio (r = 0.31, p = 1.2 x 10^-8^; Supplementary Fig. 7), an inflammation marker that is associated with increased cardiovascular risk and mortality (46, 47), but not with the proportions of CD4 T cells (r = -0.05, p = 3.8 x 10^-1^). These associations suggest that FKBP5-related inflammation could be driven by enhanced chemotaxis of granulocytes and other proinflammatory cells. As plausible mediator of this effect, we focused on interleukin-8 (IL-8), a major chemokine that recruits and activates granulocytes and other proinflammatory cells (48). Interestingly, FKBP5 downregulation has been found to suppress IL-8 production in melanoma cells (19), but no studies have examined whether FKBP5 upregulation influences IL-8 secretion by immune cells. To examine this possibility, we overexpressed *FKBP5* in Jurkat cells, a human T cell line that allowed efficient and reproducible transfection with *FKBP5* expression vectors (≈3.2-fold induction; Fig. 3B), and measured their potential to secrete IL-8. *FKBP5* overexpression nearly doubled IL-8 secretion upon immune stimulation (p = 4.4 x 10^-7^; Fig. 3C), supporting the possibility that increased *FKBP5* in T cells could drive chemotaxis of proinflammatory cells, including granulocytes.

**Figure 3.**
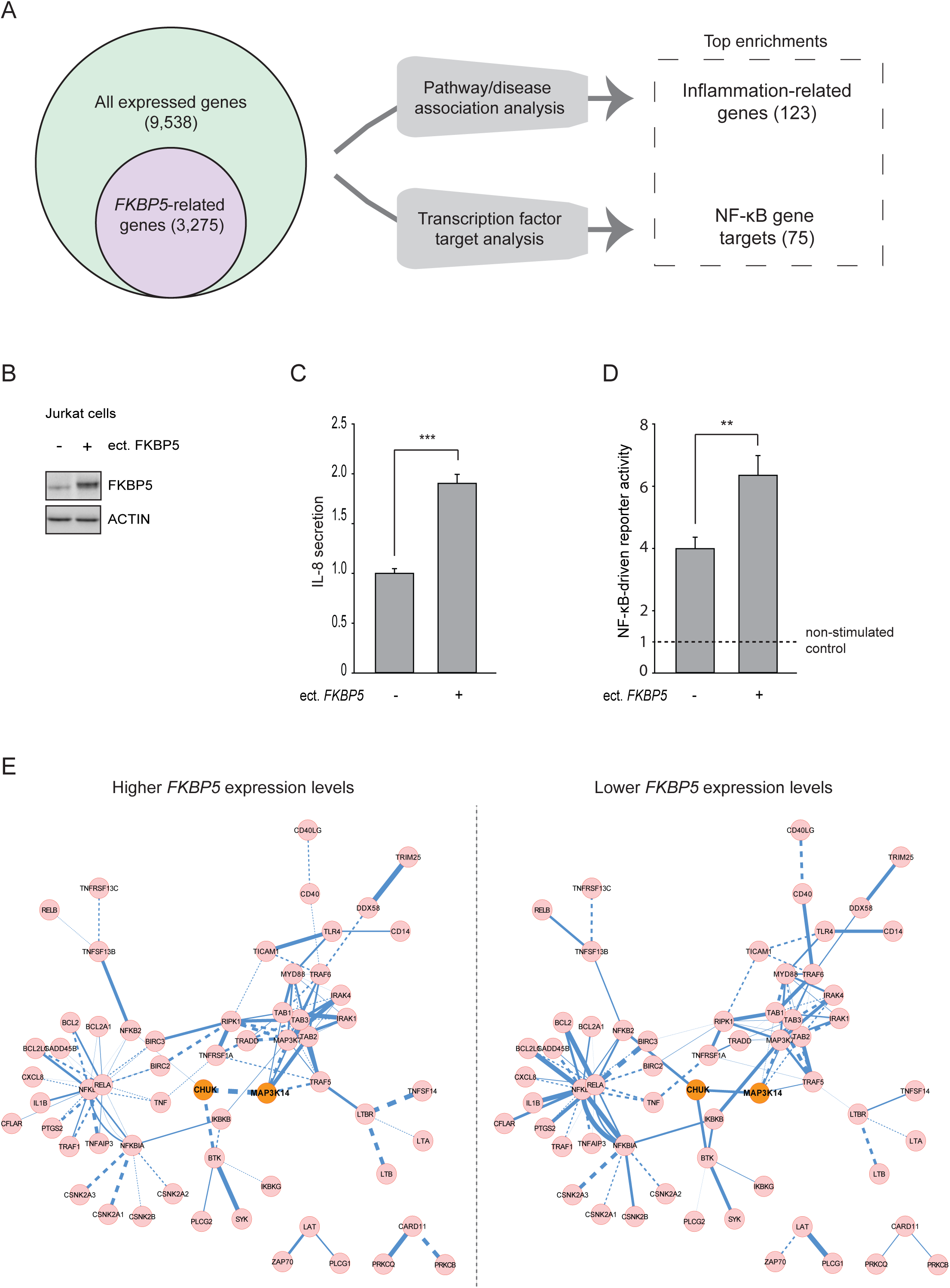
*FKBP5* upregulation promotes NF-κB-driven peripheral inflammation. **(A)** *FKBP5*-related genes in peripheral blood show enrichment for inflammation-related genes and NF-κB gene targets. Disease association and transcription factor target analyses were performed using genome-wide gene expression data in the Grady Trauma Project cohort (GTP; n = 355). The number of genes for each analysis is shown in parentheses. Statistical details are provided in Supplementary Tables 5-9. **(B)** Western blotting confirming FKBP5 overexpression in Jurkat T cells transfected with FKBP51-FLAG *vs.* cells transfected with the control vector. **(C)** *FKBP5* overexpression nearly doubles IL-8 secretion by Jurkat T cells stimulated overnight with 25 ng/ml of Phorbol-12-myristate-13-acetate and 375 ng/ml of ionomycin (PMA/I). The bar graph depicts IL-8 secretion in stimulated cell supernatants measured with ELISA from two independent experiments (t = 8.8, p = 4.4 x 10^-7^, n = 8 per condition). For each experiment, fold ratios of IL-8 secretion were calculated relative to stimulated cells expressing the control vector. IL-8 was not detectable in non-stimulated cells (not shown). **(D)** *FKBP5* overexpression increases NF-κB activity in stimulated Jurkat T cells. The bar graph depicts NF-κB reporter activity in stimulated cells measured with dual-luciferase reporter assays from three independent experiments (t = 3.2, p = 5.5 x 10^-3^, n = 9 per condition). For each experiment, fold ratios of NF-κB activity were calculated relative to non-stimulated cells expressing the control vector. **(E)** *FKBP5* expression changes are associated with extensive alterations in the NF-κB co-expression network in the GTP (n = 355). The circles depict genes encoding molecular partners of the NF-κB pathway. Continuous lines (edges) represent positive and dotted lines negative pairwise correlations corrected for expression levels of all other genes in the pathway (details in Methods). Edge widths are proportional to the absolute value of the respective correlation coefficient. The gene pair with the most robust difference in correlation between the two groups (*CHUK-MAP3K14*) is highlighted in orange. Statistical details for all gene pairs are provided in Supplementary Table 10. Error bars depict the standard error around the group mean. **\*\*** p < 10^-2^; **\*\*\*** p < 10^-3^.

To further examine whether the effects of FKBP5 on the immune system may be driven by distinct transcription factors, we performed transcription factor target analysis in the GTP cohort using the same input and reference gene sets (3,275/9,538). The strongest enrichment was observed for NF-κB (FDR-adjusted p = 4.5 x 10^-3^; Fig. 3A; Supplementary Table 8), a master immune regulator that has been linked to FKBP5 (13, 17), and this was driven by a total of 75 NF-κB gene targets (Fig. 3A; Supplementary Table 9). To experimentally confirm that FKBP5 upregulation promotes NF-κB signaling in immune cells, we performed dual-luciferase reporter assays comparing NF-κB activity between Jurkat cells overexpressing *FKBP5* and cells transfected with control vector. *FKBP5* overexpression led to increased NF-κB activity in response to immune stimulation (p = 5.5 x 10^-3^; Fig. 3D). Taken together, these findings indicate that FKBP5 upregulation in immune cells promotes NF-κB-dependent peripheral inflammation accompanied by the release of proinflammatory cytokines, such as IL8. Therefore, our subsequent analyses sought to better characterize the mechanisms through which FKBP5 impacts the NF-κB pathway.

### Changes in FKBP5 levels are associated with extensive alterations in the NF-κB co-expression network

To determine the network-level effects of FKBP5 deregulation on NF-κB signaling, we used the gene expression data in the GTP cohort (n = 355) to calculate the pairwise correlations between genes encoding molecules that directly interact within the NF-κB pathway, as defined in the KEGG Pathway Database. Each of these pairwise correlations was adjusted for the expression levels of all other genes in the pathway. These partial pairwise correlations were then compared between subjects above and those below the median split for *FKBP5* expression levels. As shown arithmetically in Supplementary Table 10 and schematically in Fig. 3E, several partial pairwise correlations within the NF-κB pathway differed between the two groups, but the strongest and only significant effect after correction for multiple comparisons was noted for the *MAP3K14*-*CHUK* pair (r_lower_ *FKBP5* = 0.13 *vs.* r_higher_ *FKBP5* = -0.28, FDR-adjusted p = 1.1 x 10^-2^, permutation p = 2.6 x 10^-3^). This effect remained robust after controlling for sex, age, cortisol, and Houseman-calculated blood cell proportions (FDR-adjusted p = 1.3 x 10^-4^, permutation p = 7.1 x 10^-3^), indicating that the effects of *FKBP5* on NF-κB signaling are not confounded by cortisol levels nor blood cell composition.

### FKBP5 upregulation promotes NF-κB signaling by strengthening the interaction of key regulatory kinases

Since FKBP5 is involved in scaffolding of regulatory protein complexes, its effects on NF-κB signaling could result from its ability to modulate protein-protein interactions between regulators of the NF-κB pathway. Intriguingly, the transcript pair most profoundly influenced by *FKBP5* levels, *MAP3K14* and *CHUK* (Fig. 3E), respectively encode the NF-kappa-B-inducing kinase (NIK) and the antagonist of nuclear factor kappa-B kinase subunit alpha (IKKα), two key regulatory kinases of the alternative NF-κB pathway. Specifically, NIK interacts with and phosphorylates IKKα at serine 176 (pIKKα^S176^), thereby activating IKKα and facilitating NF-κB signaling (49, 50).

To examine whether FKBP5 modulates the NIK-IKKα protein complex, we performed a series of protein-protein binding experiments in human Jurkat cells and peripheral blood mononuclear cells (PBMC). These experiments showed binding of FKBP5 with both NIK and IKKα and binding between NIK and IKKα (Fig. 4A). We then examined whether glucocorticoid treatment and FKBP5 upregulation can influence the FKBP5-NIK-IKKα complex. Both cell types were stimulated with DEX that robustly induces FKBP5 expression (45, 51). After confirming the DEX-induced upregulation of FKBP5 (≈2.2-fold), we found that DEX treatment significantly increased the binding between FKBP5, NIK, and IKKα in both Jurkat cells and PBMC; this increase was abolished by concomitant treatment with the recently developed selective FKBP5 antagonist SAFit1 (52) in both cell types (Fig. 4, A and B). Accordingly, these effects on protein binding were accompanied by an increase in pIKKα^S176^, whereas pIKKα^S176^ induction was abolished by treatment with SAFit1 (Fig. 4C). This effect on pIKKα^S176^ was recapitulated by *FKBP5* overexpression and again blocked by concomitant treatment with SAFit1 in Jurkat cells (Fig. 4D). Additionally, in this cell line deletion of the *FKBP5* gene with CRISPR/Cas9 abolished the effect of DEX on pIKKα^S176^ levels but did not influence vehicle-treated cells (Supplementary Fig. 8, A and B), thus mimicking the effects of SAFit1. In line with these functional effects on the NIK-IKKα complex, *FKBP5* overexpression nearly doubled NF-κB activity in Jurkat cells, whereas this effect was again prevented by concomitant treatment with SAFit1 (Fig. 4E). As schematically summarized in Fig. 4F, these convergent findings show that FKBP5 upregulation strengthens NIK-IKKα binding, increases pIKKα^S176^, and, consequently, promotes NF-κB signaling.

**Figure 4.**
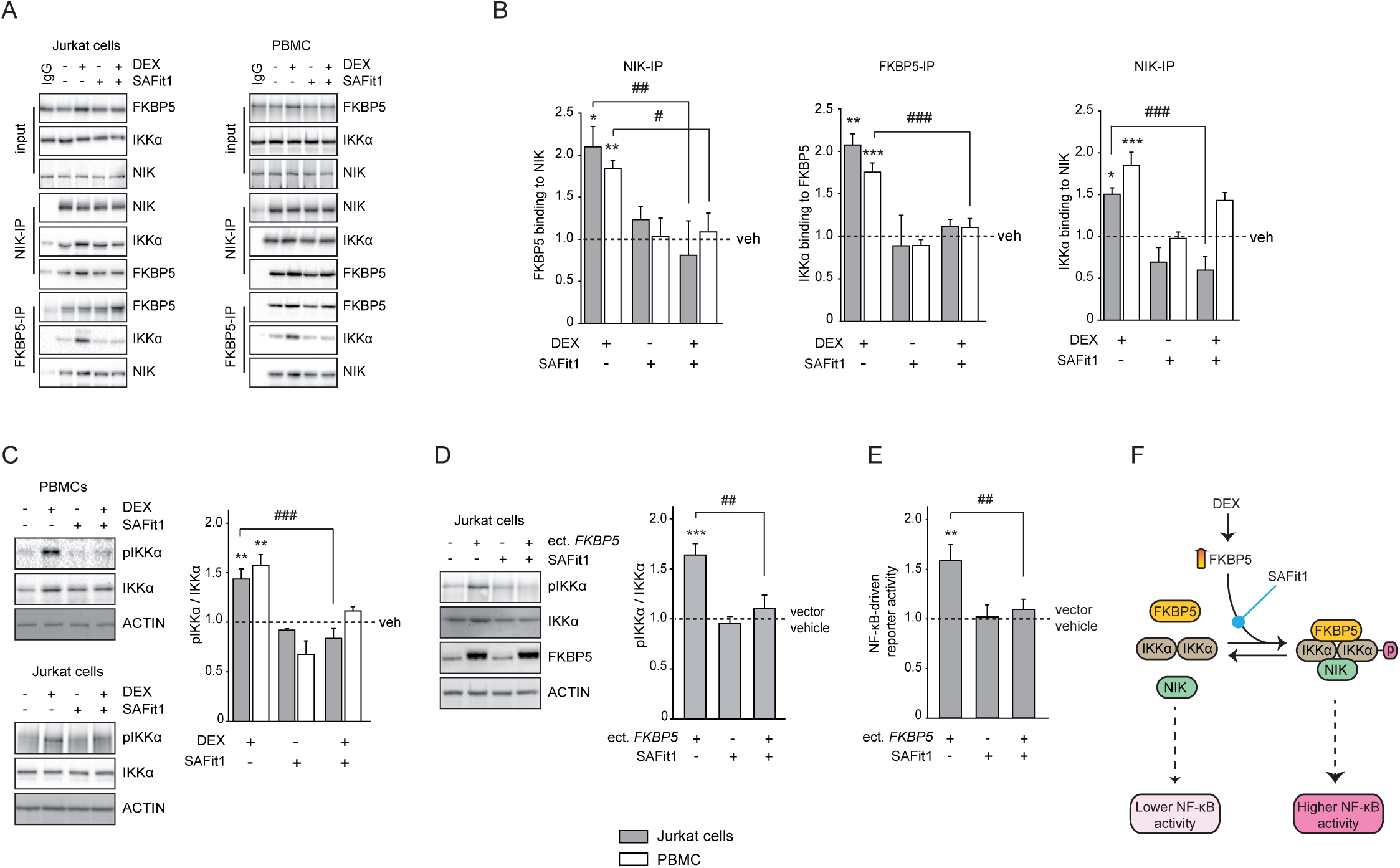
FKBP5 upregulation promotes NF-κB signaling by strengthening the binding of key regulatory kinases, and these effects are prevented by selective FKBP5 antagonism. **(A)** Immunoprecipitation (IP) for either FKBP5 or NIK followed by Western blotting in lysates from Jurkat cells or peripheral blood monocytes (PBMC) treated for 24 hours with the stress hormone (glucocorticoid) receptor agonist dexamethasone (DEX, 100nM), which robustly induces *FKBP5* expression, and/or selective FKBP5 antagonists (SAFit1, 100nM). IgG: control IP without primary antibody. **(B)** Quantifications of respective IPs showing DEX-induced increase in FKBP5-NIK-IKKα binding that is prevented by concomitant treatment with SAFit1 (Jurkat: FKBP5 to NIK binding, DEX x SAFit1 F_1,8_ = 9.3, interaction p = 1.6 x 10^-2^; IKKα to FKBP5 binding, DEX x SAFit1 F_1,8_ = 4.7, interaction p = 6.2 x 10^-2^; IKKα to NIK binding, DEX x SAFit1 F_1,8_ = 5.8, interaction p = 4.3 x 10^-2^. PBMC: FKBP5 to NIK binding, DEX x SAFit1 F_1,8_ = 5.7, interaction p = 4.4 x 10^-2^; IKKα to FKBP5 binding, DEX x SAFit1 F_1,8_ = 11.2, interaction p = 1 x 10^-2^; IKKα to NIK binding, DEX x SAFit1 F_1,8_ = 3.9, interaction p = 8.4 x 10^-2^. n = 3 biological replicates per condition). **(C)** Western blotting of Jurkat cell and PBMC lysates (n = 3 replicates per condition) showing increase in the functional phosphorylation of IKKα at serine 176 (pIKKα) by 24-hour treatment with 100nM DEX, which is prevented by 24-hour treatment with 100nM SAFit1 (Jurkat: DEX x SAFit1 F_1,8_ = 12.9, interaction p = 7 x 10^-3^; PBMC: DEX x SAFit1 F_1,8_ = 0.6, interaction p = 4.6 x 10^-1^). **(D)** Similar effects are observed when Jurkat cells are transfected with an expression construct encoding FKBP5 and treated for 24 hours with 100nM SAFit1 (ect. FKBP5 x SAFit1 F_1,12_ = 6.6, interaction p = 2.5 x 10^-2^, n = 4 replicates per condition). **(E)** *FKBP5* overexpression increases NF-κB activity in Jurkat cells stimulated overnight with 25 ng/ml of Phorbol-12-myristate-13-acetate and 375 ng/ml of ionomycin, and this increase is prevented by concomitant treatment with 100nM SAFit1 for 24 hours (ect. FKBP5 x SAFit1 F_1,32_ = 4.5, interaction p = 4.2 x 10^-2^, n = 9 replicates per condition). NF-κB reporter activity was measured with dual-luciferase reporter assays in three independent experiments. **(F)** Scheme summarizing the results from protein-protein binding and reporter gene experiments. All data are shown as fold changes compared to the control-vector vehicle-treated cells. All statistical comparisons were performed with two-way ANOVA, using either DEX treatment or *FKBP5* overexpression as the first factor and SAFit1 treatment as the second factor. Statistically significant effects were followed with Bonferroni-corrected pairwise comparisons, shown as follows:* p < 5 x 10^-2^, ** p < 10^-2^, *** p < 10^-3^, statistically significant pairwise comparisons for control *vs.* DEX or ectopic *FKBP5*; # p < 5 x 10^-2^, ## p < 10^-2^, ### p < 10^-3^, significant pairwise comparisons for vehicle *vs.* SAFit1 treatment (shown only for significant interaction terms from two-way ANOVAs). Error bars depict the standard error around the group mean.

### NF-κB signaling promotes FKBP5 expression via an NF-κB response element containing the age/stress-related CpGs

Notably, the above-identified age/stress-related *FKBP5* CpGs flank an NF-κB response element (Supplementary Fig. 9), raising the possibility that NF-κB signaling could itself modulate *FKBP5* expression in immune cells via this site. To address this possibility, we performed dual luciferase reporter gene assays using a CpG-free vector (53). We inserted into this vector the *FKBP5* sequence that surrounds the NF-κB response element and includes the two CpG sites of interest but completely lacks any other CpGs (Supplementary Fig. 9). Immune stimulation induced expression of this reporter construct in monocyte-derived human cell lines (THP-1) (Fig. 5A), thus supporting functionality of this response element in immune cells. Furthermore, *in vitro* DNA methylation of the age/stress-related *FKBP5* CpGs within this reporter construct resulted in statistically significant reduction (≈40%) of baseline expression levels and nearly abolished the induction seen with immune stimulation (Fig. 5A). To further examine whether these functional effects are mediated by alterations in NF-κB binding, we used an established biotinylated oligonucleotide-mediated chromatin immunoprecipitation (ChIP) method (54) (Fig. 5B; Supplementary Fig. 9). After confirming immune stimulation-driven NF-κB binding in THP-1 cells to the enhancer, *in vitro* DNA methylation essentially abolished NF-κB binding to the age/stress-related enhancer site (Fig. 5, C and D). Together, these findings demonstrate that NF-κB signaling –which, as we showed above, is promoted by FKBP5 (Fig. 3 and Fig. 4)-can in turn trigger *FKBP5* expression in immune cells, thereby forming a positive feedback loop that can potentiate FKBP5-NF-κB signaling. This positive feedback can thus be accentuated with decreased methylation of the NF-κB-responsive *FKBP5* enhancer, which can occur as a consequence of aging and stress.

**Figure 5.**
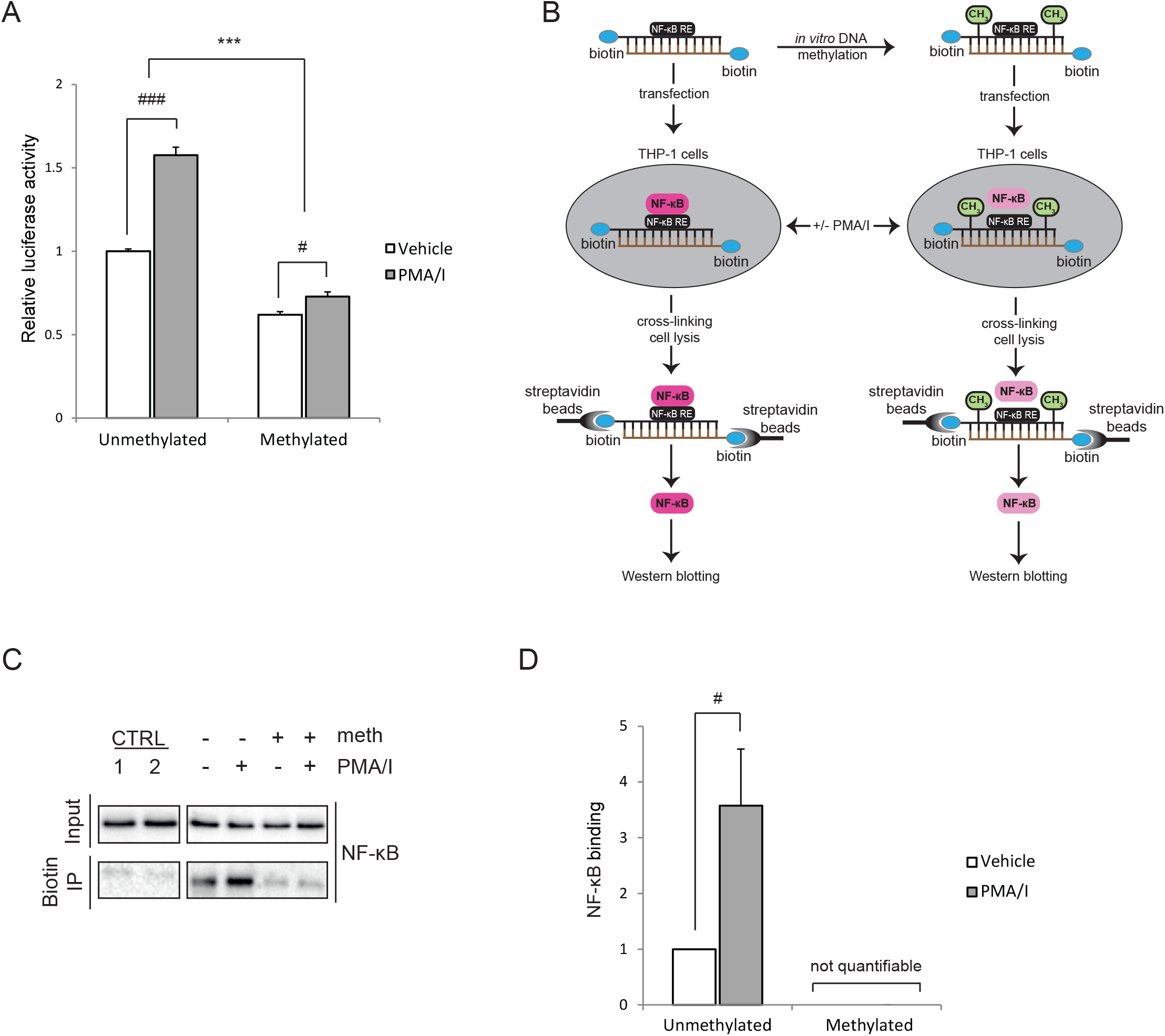
NF-κB signaling drives *FKBP5* expression via an NF-κB response element gated by the age- and stress-regulated CpGs. **(A)** Data from dual luciferase reporter gene assays using a CpG-free luciferase reporter vector to which the *FKBP5* sequence that surrounds the NF-κB response element was inserted, and which includes the two CpGs of interest but completely lacks other CpG sites (insert sequence shown in Supplementary Fig. 9). This reporter construct was *in vitro* methylated and transfected into monocyte-derived human cell lines (THP-1). Cells were then stimulated overnight with 25ng/ml Phorbol-12-myristate-13-acetate and 375ng/ml ionomycin (PMA/I), a combination that robustly induces NF-κB signaling. Data are derived from two independent experiments (n = 12 replicates per condition). Comparison was performed using two-way ANOVA with methylation and treatment as factors (F_1,44_ = 59.5, interaction p < 10^-3^), and statistically significant effects were followed with Bonferroni-corrected pairwise comparisons. **(B-D)** The effect of *in vitro* DNA methylation on PMA/I-induced NF-κB binding to the NF-κB response element was examined using biotinylated oligonucleotide-mediated chromatin immunoprecipitation (ChIP) in THP-1 cells (oligonucleotide sequence shown in Supplementary Fig. 9). Schematic summary of the experimental setup is shown in B (the lower NF-κB color intensity indicates the expected lower NF-κB binding following *in vitro* DNA methylation). After ChIP, NF-κB/p65 binding was quantified by Western blotting using antibodies specific for NF-κB (C: example blots; D: quantifications). CTRL (Control) 1: magnetic beads lacking conjugated streptavidin; CTRL 2: cells transfected with non-biotinylated oligonucleotide. Bar graph shows data derived from four independent experiments (t = 2.5, p = 4.4 x 10^-2^, n = 4 per condition). Statistical t-test compared cells carrying the unmethylated probe that were treated overnight with vehicle or PMA/I. Binding was not quantifiable for cells carrying the methylated probe. Data are always shown as fold changes compared to the vehicle-unmethylated cells. Error bars depict the standard error around the group mean. P values for pairwise comparison are shown as follows: *** p < 10^-3^, statistically significant pairwise comparisons for methylated *vs.* unmethylated. # p < 5 x 10^-2^; ### p < 10^-3^, statistically significant pairwise comparisons for vehicle *vs.* drug treatment.

### Age/stress-related decrease in FKBP5 methylation is associated with a history of acute myocardial infarction

Proinflammatory states confer risk for cardiovascular disease, most notably acute cardiovascular syndromes (55). Thus, the convergent findings presented above, indicating that lower methylation of the age/stress-related *FKBP5* CpGs derepresses *FKBP5* expression, which in turn promotes peripheral inflammation, prompted us to examine whether this lower methylation signature is also associated with higher risk for acute coronary events. To address this possibility, we used data on self-reported history of myocardial infarction (MI) that were available in both the KORA (1,648 subjects without *vs.* 62 subjects with history of MI) and the MPIP (310 controls *vs.* 8 cases) cohorts. After controlling for potential confounders (see Methods), methylation of the age-related sites was significantly lower in individuals with history of MI in both cohorts (meta-analysis p = 1.7 x 10^-2^; Fig. 6A and Supplementary Fig. 1C). This association remained significant after further adjustment for depressive phenotypes (meta-analysis p = 3.3 x 10^-2^). Given the imbalance between cases and controls, we further substantiated the identified association using permutation analysis, which yielded significant results in both cohorts (KORA, permutation p = 4.6 x 10^-2^; MPIP, permutation p = 4.8 x 10^-2^).

**Figure 6.**
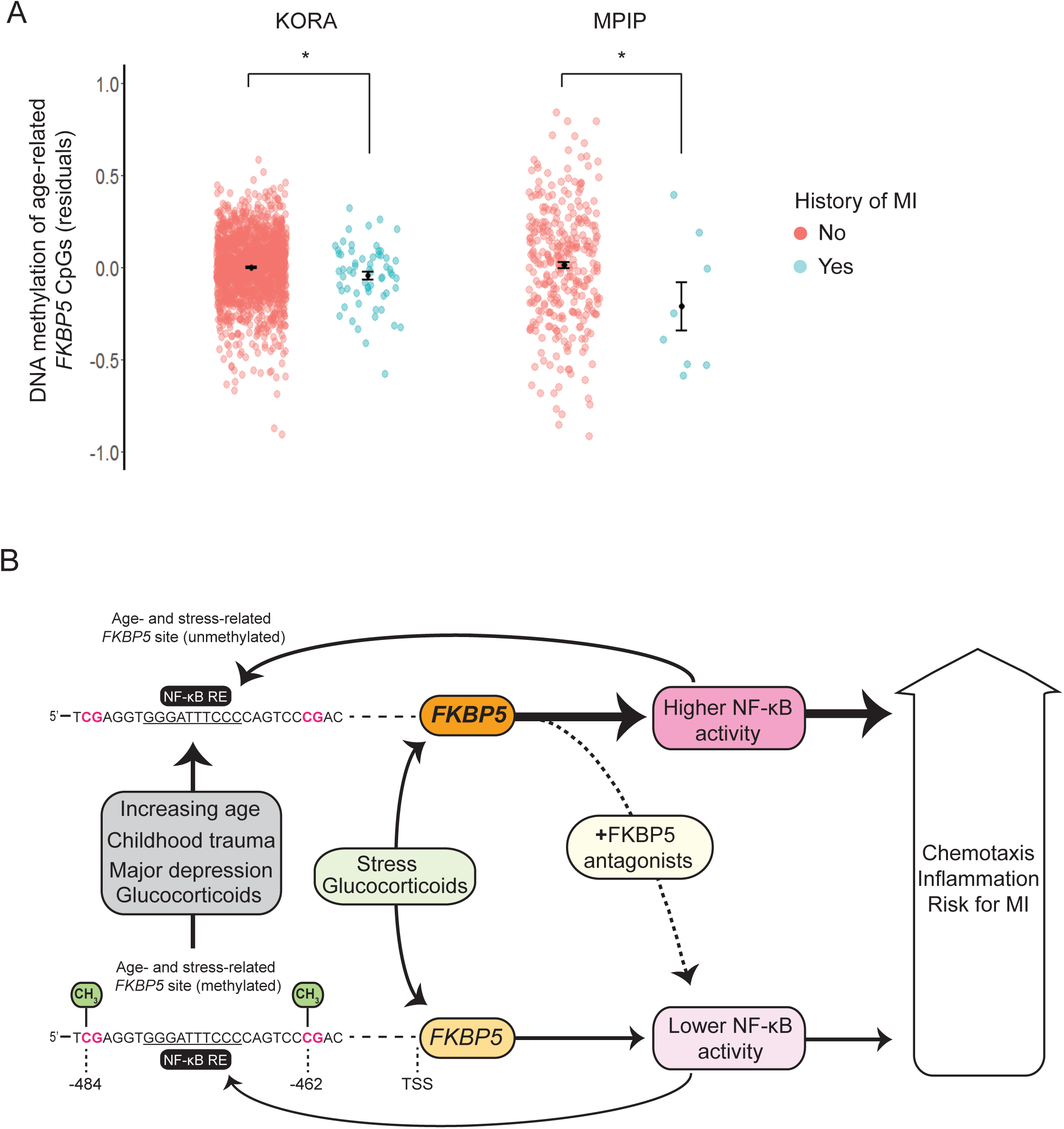
Association of age- and stress-related *FKBP5* decrease in DNA methylation with a history of myocardial infarction (MI) and overall scheme summarizing study findings. (**A**) Age- and stress-related decrease in *FKBP5* DNA methylation is associated with a history of myocardial infarction in two independent cohorts: the KORA, n = 1,648 subjects without *vs.* 62 with history of MI, β_MI_ = -0.0470, SE = 0.0231, p = 4.1 x 10^-2^, mean DNA methylation difference = 1.8%; and the MPIP, n = 310 subjects without *vs.* 8 with history of MI, β_MI_ = -0.2300, SE = 0.1177, p = 5.2 x 10^-2^, mean DNA methylation difference = 5.3%; total n = 2,028, meta-analysis p = 1.7 x 10^-2^, heterogeneity p = 1.3 x 10^-1^. The y axis depicts average DNA methylation levels of the two age-regulated *FKBP5* CpGs (cg20813374 and cg00130530), after adjusting for confounders (see Methods). Error bars depict the standard error around the group mean. **\*** p < 5 x 10^-2^. (**B**) Schematic summary of study’s findings showing how aging, childhood trauma, and depressive symptoms interact to demethylate FKBP5 at selected promoter CpGs (cg00130530 and cg20813374) located proximally (< 500 bp) upstream the transcription start site (TSS). These epigenetic changes can derepress *FKBP5* responses in immune cells, an effect that in turn promotes NF-κB signaling, whereas this is prevented in immune cells concomitantly treated with selective FKBP5 antagonists. Notably, NF-κB signaling is not only activated by FKBP5, but it can also trigger *FKBP5* transcription through an NF-κB response element that is flanked and moderated by the age/stress-related CpGs. This forms a positive feedback loop of FKBP5-NF-κB signaling that may be enhanced in individuals with lower methylation at this site. Derepressed *FKBP5* responses and NF-κB activity may promote chemotaxis of proinflammatory cells and peripheral inflammation, potentially contributing to cardiovascular risk. KORA, Cooperative Health Research in the Region of Augsburg F4 community study; MPIP, Max Planck Institute of Psychiatry depression case/control study.

## Discussion

Aging and stress-related phenotypes are associated with heightened inflammation and cardiovascular risk (5, 8-11), but the underlying molecular mechanisms remain elusive. Here we uncover a novel mechanism implicating FKBP5 in these relations. As schematically summarized in Fig. 6B, our findings suggest that aging and stress synergistically decrease DNA methylation at selected regulatory *FKBP5* CpGs that moderate the efficiency of an NF-κB-responsive enhancer. Reduced methylation at this site derepresses *FKBP5* in immune cells, an effect that promotes NF-κB-driven peripheral inflammation in part through protein-protein interactions between FKBP5 and key regulatory kinases of the NF-κB pathway. NF-κB binding to the *FKBP5* enhancer can in turn stimulate *FKBP5* expression, thereby forming a positive feedback loop of FKBP5-NF-κB signaling that potentially contributes to proinflammatory states and heightened cardiovascular risk. Notably, we find that both CRISPR-Cas9 deletion of *FKBP5* and treatment with a selective FKBP5 antagonist are able to prevent the cellular effects of stress and FKBP5 upregulation on NF-κB signaling.

By interrogating all 450K-covered CpGs spanning the *FKBP5* locus for age-related changes in DNA methylation, a biological process thought to contribute to disease states (26, 31), we identified two closely juxtaposed CpGs (separated by 22 bp) at which methylation levels are inversely associated with an interplay of aging and stress-related phenotypes. These relations were observed in both Caucasians and African Americans collectively from seven independent human cohorts, where DNA methylation was measured in whole blood. Further analyses in purified blood cells showed that age was more robustly associated with this methylation signature in CD4 T cells as compared to neutrophils, suggesting that distinct immune cell types may differ in the extent of the association of *FKBP5* methylation decrease with increasing age. These findings extend previous studies supporting the impact of both aging and life stress on the methylome (24, 25, 27-29, 56), albeit the limitations inherent to the use of human cohorts do not allow conclusive causal inferences. To offset these limitations, we experimentally tested the observed associations in an *in vitro* model of replicative senescence, wherein replicative aging and stress hormone exposure additively decreased methylation at the same age/stress-related CpGs. These convergent *in vivo* and *in vitro* observations suggest that aging and stress may together influence selected *FKBP5* CpGs across different cohorts, distinct cell types, and contexts.

Epigenetic effects involving *FKBP5* have been previously reported to occur in intronic glucocorticoid response elements, possibly as a result of glucocorticoid receptor binding to the DNA (27, 29, 57); here we identified lower methylation levels at two CpGs that include a functional NF-κB response element site and co-localize with a poised enhancer within 500 bp upstream of the *FKBP5* TSS in most immune cells, including CD4 T cells, the cell type that shows age-related decrease in *FKBP5* methylation. Taken together, this functional annotation and our associative and mechanistic data support a model whereby age/stress-related decrease in methylation at these CpGs may facilitate transcription factor binding and derepress the nearby *FKBP5* TSS in distinct immune cell types. We speculate that such decrease in methylation may additively result from the effects of cellular aging and repeated activation of the enhancer over time (age effects) coupled with glucocorticoid-induced transcriptional activation of *FKBP5* along the lifespan.

Through a combination of unbiased network analyses in human cohorts and mechanistic investigations in immune cells, we characterized a multilevel positive regulatory feedback between the stress-responsive co-chaperone FKBP5 and the NF-κB signaling cascade. More specifically, FKBP5 was found to exert pronounced effects on NF-κB-related gene networks and to promote NF-κB signaling by strengthening the interactions between NIK and IKKα, two key regulatory kinases of the NF-κB pathway. These findings are congruent with previous observations that FKBP5 downregulation can inhibit NF-κB signaling (13, 15, 17-19) and for the first time show that FKBP5 interacts with NIK and mediates the glucocorticoid-driven modulation of the NIK-IKKα regulatory complex in immune cells. Intriguingly, NF-κB can in turn trigger *FKBP5* transcription through an NF-κB response element that is flanked and moderated by the age/stress-related CpGs, thereby forming a positive feedback loop that can potentiate FKBP5-NF-κB signaling, especially in individuals with lower methylation at these *FKBP5* CpGs. Both CRISPR/Cas9 deletion of the *FKBP5* gene and treatment with the selective FKBP5 antagonist SAFit1 prevent the cellular effects of stress, as modeled *in vitro* by stress hormone treatment, and *FKBP5* overexpression on NF-κB signaling. In contrast, as shown both here and in a previous study (52), SAFit1 does not influence immune function under baseline conditions, suggesting that FKBP5 antagonism could represent a pharmaceutical intervention that —if targeted at individuals with derepressed *FKBP5*— could prevent some of the unwarranted age/stress-related alterations in immune function. However, the potential *in vivo* relevance of the pharmaceutical modulation of FKBP5-NF-κB signaling will need to be characterized by future studies.

We also find convergent evidence that FKBP5 promotes inflammation, a biological process closely linked with NF-κB signaling. This effect may in part result from the enhanced chemotaxis and recruitment of proinflammatory cells, a possibility supported by the positive association of *FKBP5* mRNA levels with the granulocyte to lymphocyte ratio and the ability of FKBP5 to augment immune cell secretion of the major chemokine, and NF-κB target, IL-8 by Jurkat cells, which is a T cell line. The latter finding extends a previous study showing that *FKBP5* downregulation suppresses NF-κB-mediated production of IL-8 in melanoma cells (19). Both IL-8 levels and the granulocyte to lymphocyte ratio are inflammatory markers associated with heightened cardiovascular risk and mortality (47, 58, 59). Together these findings suggest that older individuals with higher stress burden, who show exaggerated *FKBP5* responses, are also more likely to demonstrate heightened inflammation and acute cardiovascular risk upon stress exposure. This hypothesis is supported by our observed association, in two independent cohorts, between a history of MI and decreased *FKBP5* methylation at the age/stress-related *FKBP5* sites. *FKBP5* derepression could thus represent one molecular link underlying the previously reported association of depression and early life adversity with heightened inflammation and cardiovascular risk (3, 4, 6, 60, 61). Nevertheless, further mechanistic dissection of the potential role of FKBP5 in cardiovascular risk will require longitudinal studies examining the convergent effects of stress and aging in purified immune cell types.

In conclusion, the present study shows that aging and stress-related phenotypes are associated with lower DNA methylation at selected enhancer-related *FKBP5* sites, potentially contributing to epigenetic derepression of FKBP5 in immune cells, increased NF-κB-driven peripheral inflammation (in part through enhanced chemotaxis), and heightened cardiovascular risk. While the synergistic impact of aging and stress on disease risk is undoubtedly mediated by multiple molecular effectors and mechanisms, the present study provides novel insights by uncovering a mechanism through which stress-related phenotypes shape disease risk at the molecular level. Such insights may help to identify biomarkers for aging- and stress-related disease and may point to possible novel therapeutic interventions.

## Materials and Methods

### Human cohorts and clinical measures

The first main cohort was derived from the Grady Trauma Project (GTP), a large study conducted in Atlanta, Georgia that investigates the role of genetic and environmental factors in shaping stress responses. Participants predominantly come from an African American, urban population of low socioeconomic status (62, 63). This population is characterized by high prevalence and severity of psychosocial stress exposure and is thereby particularly relevant for examining the impact of stress-related phenotypes on genomic regulation. All African American subjects with available *FKBP5* DNA methylation and/or genome-wide gene expression data were included in the analyses. Stress-related phenotypes of interest included depressive symptoms measured by the Beck Depression Inventory (BDI) (64, 65) and childhood trauma measured with the Childhood Trauma Questionnaire (CTQ) (66). Based on a standard BDI cutoff score (64), subjects were categorized as having higher (total BDI score ≥ 19) or lower levels (total BDI score < 19) of depressive symptoms. Lifetime abuse of substances, including tobacco, alcohol, cannabis, and heroin was assessed with the Kreek-McHugh-Schluger-Kellogg scale (67). Morning serum cortisol was measured as described previously with a commercial radioimmunoassay kit (Diagnostic Systems Laboratories, Webster, TX, USA) (68). All participants provided written informed consent and all procedures were approved by the Institutional Review Boards of the Emory University School of Medicine and Grady Memorial Hospital (IRB00002114).

The second main cohort was derived from the KORA (Cooperative Health Research in the Region of Augsburg) F4 community study conducted between 2006 and 2008, a follow-up study of the fourth KORA survey S4 conducted between 1999 and 2001). Subjects were recruited from the city of Augsburg and two adjacent counties in the south of Germany (69), and DNA methylation was measured in a study subset. Depressive symptoms were assessed with the DEpression and EXhaustion subscale (DEEX scale) of the von Zerrssen symptom checklist (70). Based on a previously defined DEEX cutoff (71), subjects were categorized as having higher (total DEEX score ≥ 11) or lower levels of depressive symptoms (DEEX score < 11). Smoking was defined as current smoker, occasional smoker, former smoker, or never smoker. History of diagnosed myocardial infarction (MI) was determined using a self-reported questionnaire. The study has been conducted according to the principles expressed in the Declaration of Helsinki. Written informed consent was given by all participants. All study protocols were reviewed and approved by the local ethics committee (Bayerische Landesärztekammer).

The third main cohort comprised of Caucasian depressed and control subjects that were recruited at the Max Planck Institute of Psychiatry (MPIP). Recruitment strategies and characterization of case/control subjects have been previously described (72-74). Briefly, subjects were screened with either the Schedule for Clinical Assessment in Neuropsychiatry or the Composite International Diagnostic Screener, and diagnosis of major depressive disorder was ascertained according to the Diagnostic and Statistical Manual of Mental Disorders (DSM) IV criteria. Self-reported history of physician-diagnosed myocardial infarction was documented upon enrollment in the study. Written informed consent was obtained from all subjects, and the study was approved by the ethics committee of the Ludwig-Maximilians-University in Munich.

The impact of severe early life stress on *FKBP5* methylation was examined in a subset of the Helsinki Birth Cohort Study (HBCS) (75). The HBCS has detailed information on the separation of Finnish children from their parents, which occurred during World War II and was documented by the Finnish National Archives registry between 1939 and 1946. The subset with available DNA methylation data includes separated and non-separated (control) males. In the separated subjects group, the mean age at separation was 4.7 years (SD, 2.4 years) and the mean duration of separation 1.7 years (SD, 1 year). Based on self-report, subjects were categorized as never smokers, former smokers, occasional smokers, and active smokers. The HBCS was carried out in accordance with the Declaration of Helsinki, and the study protocol was approved by the Institutional Review Board of the National Public Health Institute. Written informed consent was obtained from all participants.

The demographics for all four main participating cohorts and relevant variables are provided in Table 1.

### DNA methylation array analyses

Genomic DNA from the GTP, the MPIP, and the HBCS cohorts was extracted from whole blood using the Gentra Puregene Blood Kit (QIAGEN). DNA quality and quantity was assessed by NanoDrop 2000 Spectrophotometer (Thermo Scientific) and Quant-iT Picogreen (Invitrogen). Genomic DNA was bisulfite converted using the Zymo EZ-96 DNA Methylation Kit (Zymo Research) and DNA methylation levels were assessed for >480,000 CpG sites using the Illumina HumanMethylation450 BeadChip (450K). Hybridization and processing was performed according to the manufacturer’s instructions as previously described(76). Quality control of methylation data, including intensity read outs, filtering (detection P value >0.01 in >50% of the samples), as well as beta and M-value calculation was done using the minfi Bioconductor R package version 1.10.2 (77). Blood cell proportions were calculated using a Minfi-based implementation of the Houseman et al. algorithm (78). We excluded X chromosome, Y chromosome, and non-specific binding probes (79), as well as probes if single nucleotide polymorphisms (SNPs) were documented in the interval for which the Illumina probe is designed to hybridize. Given that the GTP cohort includes individuals from different ethnicities, we also removed probes if they were located close (10 bp from query site) to a SNP which had Minor Allele Frequency of ≥0.05, as reported in the 1,000 Genomes Project, for any of the populations represented in the samples. All data were normalized with functional normalization (FunNorm) (80), an extension of quantile normalization included in the R package *minfi.* Technical batch effects were identified by inspecting the association of the first principal component of the methylation levels with bisulfite conversion plate, plate position, array, slide, slide row and slide column, and by further visual inspection of principal component plots using the shinyMethyl Bioconductor R package version 0.99.3 (81). Batch effect removal was then performed with the Combat procedure as implemented in the sva package (82), and the batch-corrected data were used for further analyses. The raw methylation data for the GTP cohort have been deposited into NCBI GEO (GSE72680).

For the KORA study, genomic DNA (1 µg) from 1,814 samples was bisulfite converted using the EZ-96 DNA Methylation Kit (Zymo Research, Orange, CA, USA) according to the manufacturer’s protocol, with the incubation conditions recommended for the Illumina Infinium Methylation Assay. Raw methylation data were generated by BeadArray Reader and extracted by GenomeStudio (version 2011.1) with methylation module (version 1.9.0). Data were preprocessed using R version 3.0.1 (http://www.r-project.org/) (83). Probes with signals from less than three functional beads and probes with a detection p-value > 0.01 were defined as low-confidence probes. As probe binding might be affected by SNPs in the binding area, cytosine-guanine dinucleotides sites (CpGs) in close proximity (50bp) to SNPs with a minor allele frequency of at least 5% were excluded from the dataset. Color bias adjustment using smooth quantile normalization method as well as background level correction based on negative-control probes present on the Infinium HumanMethylation BeadChip was performed for each chip using the R package lumi (version 2.12.0) (84). Beta values corresponding to low-confidence probes were then set to missing, and samples as well as CpGs were subjected to a 95% detection rate threshold, where samples and CpGs with more than 5% low-confidence probes were removed from the analysis. Finally, beta-mixture quantile normalization (BMIQ) was applied to correct the shift in the distribution of the beta values of the InfI and InfII probes (85). BMIQ was done using the R package wateRmelon (version 1.0.3) (86). The KORA F4 samples were processed on 20 96-well plates in 9 batches; technical batch effects were investigated through principle component and eigen-R2 analysis based on positive control probes (87). Batches were then corrected by including the first principle components as covariates in all regression models.

DNA methylation analyses included 45 CpGs covered by the 450K that are located within or in close proximity (10kb upstream or downstream) to the *FKBP5* locus. All 45 CpGs were measured in the KORA, whereas one CpG (cg00052684) did not pass quality control in the other cohorts. In all cohorts, we controlled for the potential confounding effect of cell type heterogeneity with a DNA methylation-based approach (78) that employs 450K array data to calculate blood cell proportions and that is widely used in most of the large epigenome-wide and other association studies (88, 89) and has repeatedly been validated (90, 91). The calculated blood cell proportions were included as covariates in all DNA methylation analyses. To further rule out the confounding effect of distinctive cell type patterns, we also analyzed DNA methylation data from FACS-sorted CD4 T cells and neutrophils that are publicly available in NCBI GEO (GSE67705) (35). Lastly, we used an additional dataset of male and female subjects (n = 213) with both 450K data and differential blood counts available at the MPIP to rule out potential confounding by counted blood cell types. Analytical approaches in this dataset were performed as previously described (24).

### Functional annotation of DNA methylation sites

Annotation of the identified CpGs was performed using the Roadmap Epigenome Browser (http://epigenomegateway.wustl.edu/browser/roadmap/) for all available tracks (H3K4me1, H3K4me3, H3K27ac, H3K27me3, H3K9me3, H3K36me3, and methylC-Seq) and for the genetic location surrounding the age/stress-related sites (hg19, chr6:35654000-35660000). To assess whether the two sites co-localize with specific chromatin states, we used the 15-state ChromHMM annotation of the Roadmap Epigenomics project and calculated the position-based overlap of all 15 states among the 127 available epigenomes (42).

### Gene expression arrays

Genome-wide gene expression data were measured in 355 African American subjects from the GTP. Whole blood RNA was collected, processed, and hybridized to Illumina HumanHT-12 v3 and v4 Expression BeadChips (Illumina, San Diego, CA, USA) as previously described (45, 76). The raw microarray scan files were exported using the Illumina Beadstudio program 13 and further analyzed in R (<u>www.R-project.org</u>). Microarray data were transformed and normalized via the variance stabilizing normalization with the use of Illumina internal controls (92). Empirical Bayes method was used to control for potential confounding as a result of batch effects (82). Six pairs of technical replicates were used to confirm data reproducibility (average Pearson correlation 0.996). The raw gene expression array data for the GTP study have been deposited to GEO (GSE58137).

### Population stratification

To control for potential confounding by population stratification, we used genome-wide SNP data. In the GTP cohort, of the 700 k SNPs present on the Omni Quad and Omni express arrays, 645,8315 autosomal SNPs were left after filtering with the following criteria: minor allele frequency of >1 %; Hardy-Weinberg equilibrium of 0.000001; and genotyping rate of >98 %. The MPIP cohort was genotyped using the Illumina 300k, 610k, and Omni express arrays. For each chip array, quality control was performed separately following the same quality control protocol like in the GTP. After QC, we used the overlap of 168,138 SNPs across all chip types. The samples were clustered to calculate rates of identity by descent (IBD). We then ran multidimensional scaling analysis on the IBD matrix using PLINK2 (https://www.cog-genomics.org/plink2) and plotted the first ten axes of variation against each other. No outliers were detected. The first two principal components were used as covariates in regression models to adjust for population stratification.

### Pathway analyses

*FKBP5* mRNA levels were correlated with the expression of all genes with at least one array probe detected above background in peripheral blood in the GTP cohort. For genes with more than one probes, gene (mRNA) expression levels were calculated by averaging their levels. Using the set of FDR-corrected genes correlating with *FKBP5* as input and the set of genes expressed above background as reference, we then implemented disease association and transcription factor target analysis using the WEB-based GEne SeT AnaLysis Toolkit (WebGestalt 2013; http://www.webgestalt.org/webgestalt_2013/) (93, 94). Enrichment analysis for disease-associated genes is based on GLAD4U as the source database. Enrichment analysis for the targets of transcription factors is performed using MSigDB as the source database. Both analyses were performed with a hypergeometric test, whereby the minimum number of genes for the enrichment analysis was set at 5. Both analyses were FDR-corrected for multiple testing.

For analysis involving the co-expression network of the NF-κB (nuclear factor kappa-light-chain-enhancer of activated B cells), the list of NF-κB-related genes was acquired from the KEGG Pathway Database (http://www.genome.jp/dbget-bin/www_bget?pathway:hsa04064). Using the gene expression array data in the GTP as described above, the pairwise correlation coefficients between gene pairs encoding molecules that directly interact along the NF-κB pathway were calculated and adjusted for the expression levels of all other pathway partners using the R package GeneNet (95). The adjustment is made by Gaussian Graphical Model (GGM), which measures the degree of association between two genes by controlling for the expression of other genes. For genes involved in the same pathway, the correlation of every two genes can be directed and undirected. For two genes of interest, using the Pearson correlation coefficient will give a misleading result if they are related to another gene(s). A partial correlation coefficient (pcor) measures the degree of association between two random variables after controlling for the effect of a set of random variables. Suppose there are three genes x, y, and z, the pcor of gene x and gene y conditioning on effect of gene z is given as:

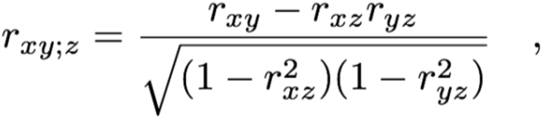

where r_xy_, r_xz_, and r_yz_ are the standard Pearson product-moment correlation coefficients. Let G = {g_1,…,g_m} be the expression matrix of m genes in a pathway. A pathway-based pcor denotes the pairwise correlation of gene x and gene y with the effect of the remaining m-2 genes in same pathway removed, which is calculated by:

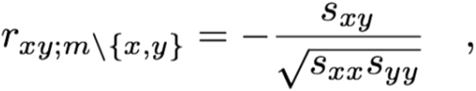

where s_xy_ is the (x,y) element of the inverse of the correlation matrix. The effect of age, sex, cortisol, and Houseman-calculated blood cell proportions is removed by including the respective variables into G.

These partial pairwise correlations were then compared between subjects with higher and those with lower *FKBP5* expression as defined by a median split of *FKBP5* mRNA levels. To assess the significance of the difference between two correlation coefficients, the coefficients have to be converted to z values using Fisher's z transformation (96). The formula for the transformation is:

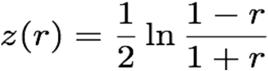

where ln is the natural logarithm. For sample sizes of n1 and n2 find the z of the difference between the z transformed correlations divided by the standard error of the difference of two z scores (https://CRAN.R-project.org/package=psych Version = 1.7.8.):

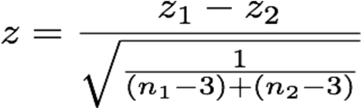

Then z is approximately normally distributed with a mean of 0 and a standard deviation of 1. In order to account for multiple testing, we applied FDR correction to obtain a corrected estimate of the significance level. Lastly, to test the robustness of the change in partial correlations between the two groups, the *FKBP5* high/low group assignments for each pair were permuted 10,000 times across samples.

### Cell culture

Cell culture experiments involving immune cells were conducted in peripheral blood mononuclear cells (PBMC), Jurkat cell lines (ATCC, TIB-152), or THP-1 (ATCC, TIB-202). For PBMC isolation, the whole blood of healthy volunteers was collected via venipuncture, diluted with PBS, carefully loaded on Biocoll solution (BioChrom, L6113), and centrifuged at 800 g for 20 min without brake. PBMC were enriched by selecting the interphase of the Biocoll gradient and were then washed two times with ice-cold PBS and resuspended in medium. All cell types were maintained in RPMI (Gibco) supplemented with 10% FCS and 1% Antibiotic/Antimycotic (Thermo Fisher scientific Inc., Schwerte, Germany). *FKBP5* overexpression in Jurkat cells was performed using a previously described FKBP51-FLAG (97). For all drug treatments of immune cells, cells were left after seeding to rest overnight and were subsequently incubated overnight with vehicle (0.05% DMSO), 25ng/ml Phorbol-12-myristate-13-acetate (PMA, Sigma, P1585) and 375ng/ml ionomycin (Sigma, I0634), 100 nM dexamethasone (DEX; Sigma, D4902), and/or 100 nM SAFit1 as indicated per experiment. Cell transfections with amounts of 2 µg of plasmid DNA were performed using the Amaxa Nucleofector Device and the Cell Line Nucleofector Kit V (Lonza, Basel, Switzerland).

To model replicative senescence *in vitro*, IMR-90 cells (I90-83, Passage 4) were obtained from the Coriell Institute for Medical Research (Camden, New Jersey, USA) and grown in DMEM medium (41966029, Thermo Fisher scientific Inc., Schwerte, Germany) supplemented with minimum essential medium non-essential amino acids (Thermo Fisher scientific Inc., Schwerte, Germany), 15 % FCS (Thermo Fisher scientific Inc., Schwerte, Germany) and 1 % Antibiotic/Antimycotic in an incubator under 37 °C and 5 % CO2 conditions. Cells were split at a confluency of 80-90% after a wash step with DPBS (Thermo Fisher scientific Inc., Schwerte, Germany) using Trypsin-EDTA (Thermo Fisher scientific Inc., Schwerte, Germany) (8-10 min at 37°C). Cells were aged in culture, and replicative age was defined according to the population doubling level (PDL) as previously described(39) as either young (PDL = 22) or old (PDL = 42). To model stress *in vitro*, cells of young and old age (n = 4 replicates each) were split and treated in parallel with either 100nM DEX or vehicle (DMSO) for 7 days. For DNA extraction, cells were pelleted and extracted using the NucleoSpin^®^ Tissue Kit (Macherey-Nagel GmbH & Co.KG, Dueren, GER) following the manufacturer’s instructions.

### Targeted bisulfite sequencing

To validate the age-related sites using a non-hybridization-based DNA methylation method, we performed targeted bisulfite sequencing using the Illumina MiSeq, 200 ng to 500 ng of genomic DNA was bisulfite treated using the EZ DNA Methylation Kit (Irvine, California). Target enrichment was achieved by PCR amplification using 20 ng of bisulfite converted DNA, bisulfite specific primer (FKBP5_TSS_5.1_FW:TTTTTGTTTTTTGTGGGGGT; FKBP5_TSS_5.1_RW: AACCTCTTCCTATTTTAATCTC), Takara EpiTaq HS Polymerase (Saint-Germain-en-Laye, France), and the following cycling conditions: 94°C – 3 min, 2x (94°C – 30s, 62.1°C – 30s, 72°C – 30s), 5x (94°C – 30s, 60.1°C – 30s, 72°C – 30s), 8x (94°C – 30s, 58.1°C – 30s, 72°C – 30s), 34x (94°C – 30s, 56.1°C – 30s, 72°C – 30s), 72°C – 5 min, 4°C – ∞. Equimolar pooled amplicons of each sample were cleaned up by a double size selection (200-500 bp) using Ampure XP beads (Krefeld, Germany). The Illumina TruSeq DNA PCR-Free HT Library Prep Kit (San Diego, California) was used for library generation according to the manufacturer’s instructions. Paired-end sequencing was performed on a Illumina MiSeq Instrument (San Diego, California) with the MiSeq Reagent Kit v3 (2x 300-cycles) with the addition of 30% of PhiX Library. The quality of sequencing reads was performed with FastQC (http://www.bioinformatics.babraham.ac.uk/projects/fastqc). After trimming (Cutadapt v1.9.1) Illumina adapter sequences, Bismark v.0.15.0 was used for the alignment to a restricted reference that was based on the PCR targets. An in-house Perl script was used to stitch paired-end reads and remove low quality ends of paired-end reads if they overlapped. The methylation levels for all CpGs, CHGs, and CHHs were quantified using the R package methylKit. Subsequently, a three-step quality control was utilized to validate the methylation calls. First, PCR artifacts introducing CpGs at 0 or 100% methylation level were removed. Second, CHH methylation levels were analyzed, and samples with insufficient bisulfite conversion rate (< 95%) were removed. Finally, CpG sites with coverage lower than 1000 reads were excluded.

For DNA methylation measurements in IMR-90 cells, *FKBP5* methylation of CpG sites corresponding to 450K cg20813384 and cg00130530 (Chr6: 35657180 & Chr6:35657202, respectively; Assembly: hg19) was assessed by targeted bisulfite pyrosequencing in four biological replicates. For each replicate, 500 ng genomic DNA was bisulfite converted using the EZ DNA Methylation Kit (Zymo Research Corporation, CA, USA). Bisulfite converted DNA was amplified in a 50 µl reaction mix (1 µl DNA; each bisulfite specific Primer with a final concentration of 0.2 µM, FKBP5_TSS_5_F: TTTTTGTTTTTTGTGGGGGT & FKBP5_TSS5_R_biot: biotin-AACCTCTTCCTATTTTAATCTC) using the Takara EpiTaq HS Polymerase (Clontech, Saint-Germain-en-Laye, France). Cycling conditions of the touchdown PCR were 94°C for 3 min, 2x (94°C - 30s, 62.1 °C - 30s, 72°C - 30s), 5x (94°C - 30s, 60.1 °C - 30s, 72°C - 30s), 8x (94°C - 30s, 58.1 °C - 30s, 72°C - 30s), 34x (94°C - 30s, 56.1 °C - 30s, 72°C - 30s), 72°C for 5 min, and cooling to 4°C. Pretreatment of PCR amplicons was facilitated with the PyroMark Q96 Vacuum Workstation (QIAGEN GmbH, Hilden). Sequencing of FKBP5 CpGs was performed on a PyroMark Q96 ID system (QIAGEN GmbH, Hilden) using PyroMark Gold Q96 reagents and the following sequencing Primer: FKBP5_TSS_5_Seq1 (CpG 35657180, 35657202 / hg19): AATTTTATTAAGTTTAAGATGTTTA. The PyroMark Q96 ID Software 2.5 (QIAGEN GmbH, Hilden) was used for data analysis.

### Generation of FKBP5 knockout Jurkat cells

Jurkat cells were transfected with a pool of three clustered regularly interspaced short palindromic repeats-associated Cas 9 (CRISPR/Cas9) plasmids containing gRNA that targets human *FKBP5* and a GFP reporter (Santa Cruz, sc-401560). 36 hours post transfection, cells were FACS-sorted for GFP as single cells into a 96-well plate using BD FACSARIA III. Single clones were expanded and Western blotting was used to confirm the successful knockout of *FKBP5*.

### Co-immunoprecipitation experiments (CoIPs)

CoIPs of endogenous or FLAG-tagged FKBP5 with endogenous IKKα and NIK were performed in Jurkat cells and PBMC using previously described methods (98). 5 x 106 cells were transfected with 2 µg of the respective expression plasmids. After 3 days of cultivation in medium, cells were lysed in CoIP buffer containing 20 mM Tris-HCl (pH 8.0), 100 mM NaCl, 1 mM EDTA, and 0.5% Igepal, complemented with protease inhibitor cocktail (Roche). This was followed by incubation on an overhead shaker for 20 min at 4°C. The lysate was cleared by centrifugation, the protein concentration was determined by performing a BCA assay, and 1.2 mg of lysate was incubated with 2.5 µg of FLAG antibody overnight at 4°C. 20 µl of BSA-blocked Protein G Dynabeads (Invitrogen, 100-03D) were added to the lysate-antibody mix, followed by 3 h of incubation at 4°C. The beads were washed three times with PBS, and protein-antibody complexes were eluted with 100 µl of 1 x FLAG-peptide solution (Sigma, 100–200 µg/ml, F3290) in CoIP buffer for 30 min at 4°C. 5–15 µg of the cell lysates or 2.5 µl of the immunoprecipitates was separated by SDS-PAGE and electrotransferred onto nitrocellulose membranes (Western blotting). Opto-densimetric determination of immunodetected bands was carried out on ChemiDoc MP (BioRad). Signals of co-immunoprecipitated proteins were normalized to signals of input-corrected immunoprecipitated proteins.

### In vitro DNA methylation

*In vitro* DNA methylation of biotinylated probes and reporter constructs (detailed sequences in Supplementary Fig. 9) was carried out by *Sss*I CpG Methyltransferase (New England Biolabs, M0226S) according to the manufacturer’s instructions. Successful methylation was confirmed by restriction digest with *Xho*I, whereby linearization of the plasmids is blocked by CpG methylation. Restriction digests were controlled by agarose gel electrophoresis. Methylated and unmethylated constructs were subsequently transfected into THP-1 cells as described below. Controls underwent the same procedure except that enzyme and SAM was not added to the reaction.

### Biotinylated oligonucleotide-mediated chromatin immunoprecipitation (ChIP)

Biotinylated oligonucleotide-mediated ChIP was performed in THP-1 cells using a previously established method(54) (schematically outlined in Fig. 5B). Briefly, single-stranded complementary biotinylated oligonucleotide probes (length 70 bp) including the DNA methylation sites and response element of interest (Supplementary Fig. 9) were synthesized (Eurofins Genomics, Ebersberg, Germany), and after annealing underwent *in vitro* DNA methylation as described above. Subsequently, 0.5 pmol/106 cells of either the methylated or unmethylated probes were transfected into THP-1 cells (1.5-2 x 106 cells per replicate). After an overnight rest, the cells were incubated for 24 hours at 37°C with either vehicle (DMSO) or PMA/I (concentrations as above), a stimulus that robustly induces NF-κB signaling. Cells subsequently underwent cross-linking with 1% formaldehyde at room temperature for 15% and were then lysed in RIPA buffer (Merck, 20-188, completed with protease inhibitor cocktail). Lysates were immunoprecipitated using streptavidin-coupled magnetic beads (Dynabeads M-280, Thermo Scientific, 11205D) or control beads (Protein G Dynabeads, Thermo Scientific, 10007D), and both input and eluates were quantified for NF-κB/p65 binding by Western blotting.

### Western Blot Analysis

Unless otherwise indicated, protein extracts were obtained by lysing cells in 62.5 mM Tris, 2% SDS, and 10% sucrose, supplemented with protease (Sigma, P2714) and phosphatase (Roche, 04906837001) inhibitors. Samples were sonicated and heated at 95°C for 5 min. Proteins were separated by SDS-PAGE and electro-transferred onto nitrocellulose membranes. Blots were placed in Tris-buffered saline (TBS; 50 mM Tris-Cl, pH 7.6; 150 mM NaCl), supplemented with 0.05% Tween (Sigma, P2287) and 5% non-fat milk for 1 h at room temperature and then incubated with primary antibody (diluted in TBS/0.05% Tween) overnight at 4°C. The following primary antibodies were used: FLAG (1:7,000, Rockland, 600-401-383), FKBP5/FKBP51 (1:1,000, Bethyl, A301-430A; 1:1000, Cell Signaling, #8245), NF-κB/p65 (1:1000, Cell Signaling, #8242), IKKα (1:1000, Cell Signaling, #2682), pIKKα^S176^ (1:1000, Cell Signaling, #2078), NIK (1:1000, Cell Signaling, #4994), and Actin (1:5,000, Santa Cruz, sc-1616). Subsequently, the blots were washed and probed with the respective horseradish-peroxidase or fluorophore-conjugated secondary antibody for 2 h at room temperature. The immuno-reactive bands were visualized either by using ECL detection reagent (Millipore, WBKL0500) or directly by excitation of the respective fluorophore. Recording of the band intensities was performed with the ChemiDoc MP system from Bio-Rad. All protein data were normalized to Actin, which was detected on the same blot in the same lane (multiplexing). In the case of IKKα phosphorylation and to rule out confounding by changes in total IKKα levels, we normalized pIKKα by calculating its ratio to total IKKα. We obtained indistinguishable results when normalizing pIKKα to Actin.

### Dual-luciferase reporter gene assays

The functional effect of differential methylation in the age/stress-related *FKBP5* site was analyzed using a CpG-free luciferase reporter construct (53). The genomic region of interest (length 224 bp; Supplementary Fig. 9) was synthesized and inserted into the *Spe*I and *Pst*I sites of the CpG-free vector plasmid (Eurofins Genomics, Ebersberg, Germany). All constructs were verified by sequencing. THP-1 cells were transfected with either methylated or unmethylated reporter plasmid (800 ng/106 cells) and with the previously described Gaussia-KDEL control plasmid (100 ng/106 cells) (99). To control for differences in transfection efficiency, the NF-κB-driven reporter gene activity was calculated as the ratio of firefly (*Photinus Pyralis*) to *Gaussia* luciferase signals.

To assess the impact of *FKBP5* overexpression on NF-κB activity, 1 x 106 Jurkat cells were transfected with NF-κB luciferase reporter (1µg, Promega, E8491) and Renilla (300 ng, Promega, E6921) control plasmids, as well as with either FKBP5-FLAG (1 µg) or control vector (pRK5; 1 µg). Immediately after transfection, cells were seeded on 96-well plates at a density of 20,000 cells/well and were left to rest overnight. On the next day, cells were incubated for 2 hours with DMSO vehicle or 100nM SAFit1 and were then stimulated overnight with PMA/I (concentration as above). Cells were then lysed in lysis buffer (Promega, E1941) and Firefly and Renilla luciferase activities were recorded with the TriStar^2^ S LB 942 microplate reader (Berthold, Bad Wildbad, Germany) following a previously described protocol (100). To control for differences in transfection efficiency, the NF-κB-driven reporter gene activity was calculated as the ratio of firefly to *Renilla* luciferase signals. For each experiment, fold ratios of NF-κB activity were calculated by comparison to cells expressing the control vector.

### Enzyme-linked immunosorbent assay (ELISA) for human interleukin-8 (IL-8)

1 × 106 Jurkat cells were transfected with either FKBP5-FLAG (1 µg) or control vector (1 µg), were seeded in 24-well plates at a density of 500,000 cells/well and, after overnight rest, were stimulated with PMA/I (concentration as above). Supernatants were collected the next day, cleared by centrifugation at 125g, and stored at -80°C until IL-8 measurement with ELISA, which was performed with a commercially available kit (Merck Millipore, EZHIL8) according to the manufacturer’s instructions.

### Statistics

All statistical analyses involving DNA methylation used M-values, which are suggested to show superior statistical performance as compared to Beta-values (101). To average the methylation levels of the two age-related *FKBP5* CpGs, we calculated the mean of the respective Beta-values and then transformed each mean to the corresponding M-values as previously suggested (101). All p values reporting the statistical significance of DNA methylation analyses originate from tests using M values, and the residuals from the respective regression models are shown on the main figures; however, to more intuitively depict methylation results, Supplementary figures also show the DNA methylation Beta-values for selected panels.

Linear regression models examined the effect of age and stress-related phenotypes on DNA methylation, while including as covariates all the potential confounders that were available in the respective cohorts. Covariates used in each cohort were as follows: age, sex, blood cell proportions, the first two genome-wide SNP-based principal components, and smoking and other substance use in the GTP cohort; age, sex, blood cell proportions, smoking status, and the first two Eigen-R2-derived principal components in the KORA cohort; age, sex, case/control status, blood cell proportions, and the first two genome-wide SNP-based principal components in the MPIP cohort; and age, smoking status, and blood cell proportions in the HBCS cohort. The model validating the association of age with the age-related CpGs using targeted bisulfite sequencing in a small sample of female subjects was adjusted for smoking and blood cell proportions. Linear regression models examining *FKBP5* expression in the GTP cohort as the dependent variable of interest included as covariates age, sex, the first two SNP-based principal components, and blood cell proportions; these models tested methylation of the age-related *FKBP5* CpGs, age, stress-related phenotypes, and cortisol as the independent variables of interest. Lastly, linear regression models examined the association of history for MI with lower methylation of the age-related CpGs in the KORA and MPIP cohorts, while controlling for age, sex, blood cell proportions, and the first two principle components. For stratified analyses in the GTP cohort, we performed median splits of the respective continuous variables; this was performed for methylation levels of the age-related *FKBP5* CpGs to distinguish subjects with higher vs. lower methylation, for CTQ scores to stratify individuals in high-vs. low-trauma groups, for chronological age to stratify younger vs. older individuals, and for *FKBP5* mRNA levels to distinguish subjects with higher vs. lower *FKBP5* expression. All p values reporting the statistical significance of regression models are after correction for relevant covariates as described above.

Experimental data were normally distributed and were tested using student’s t-test when comparing two groups and with two-way analysis of variance (ANOVA) when examining two factors of interest. Significant effects in the two-way ANOVA were followed with Bonferroni-corrected pairwise comparisons.

Experimental data were analyzed in Sigma Plot version 13.0 (Systat Software Inc.). All other statistical tests were performed in R version 3.1.0 (http://www.r-project.org/) (83). All meta-analyses were performed using the package *rmeta* in R, which combines the regression coefficients and standard errors from individual cohorts. The level of statistical significance was set a priori at 0.05 (5 × 10^-2^). All reported p values are two-tailed and nominal, unless corrected for multiple testing or permutation-based as indicated and reported in the manuscript.

## Supporting information

## Acknowledgements

This work was supported by a Marie-Sklodowska Curie fellowship (H2020 grant# 653240) to ASZ, a grant from the National Institute of Mental Health (MH071538) to KJR, a European Research Council starting grant within the FP7 framework to EBB (grant# 281338, GxE molmech), a grant from the National Institute of Mental Health (U19 MH069056), a grant by the German Federal Ministry of Education and Research (BMBF) through the Integrated Network IntegraMent (Integrated Understanding of Causes and Mechanisms in Mental Disorders), under the auspices of the e:Med Programme (grant # 01ZX1314J), to EBB, and by the Academy of Finland (grant numbers 284859, 2848591, 312670).

## Author contributions

ASZ and EBB conceived and designed the study. ASZ, TR, NCG, and EBB designed and interpreted wet lab experiments. ASZ, KH, MK, JCP, SM, AH, TW, MR, and NCG performed wet lab experiments. ASZ, MJ, JA, SR, and BMM designed and performed pathway analyses. TCR, BB, KJR, AKS, and EBB contributed DNA methylation, gene expression, and phenotypic data from the GTP cohort. JB, RTE, MW, SW, SK, CG, and KHL contributed DNA methylation and phenotypic data from the KORA F4 community cohort. ASZ, TCR, SI, SL, and EBB contributed DNA methylation and phenotypic data from the MPIP cohort. JL, KR, JGE, and AJD contributed DNA methylation and phenotypic data from the HSBC cohort. FH provided the FKBP5 antagonist. ASZ wrote the manuscript, with input from EBB. All authors commented and approved the final version of the manuscript.

---

**Supplementary Figure 1.**
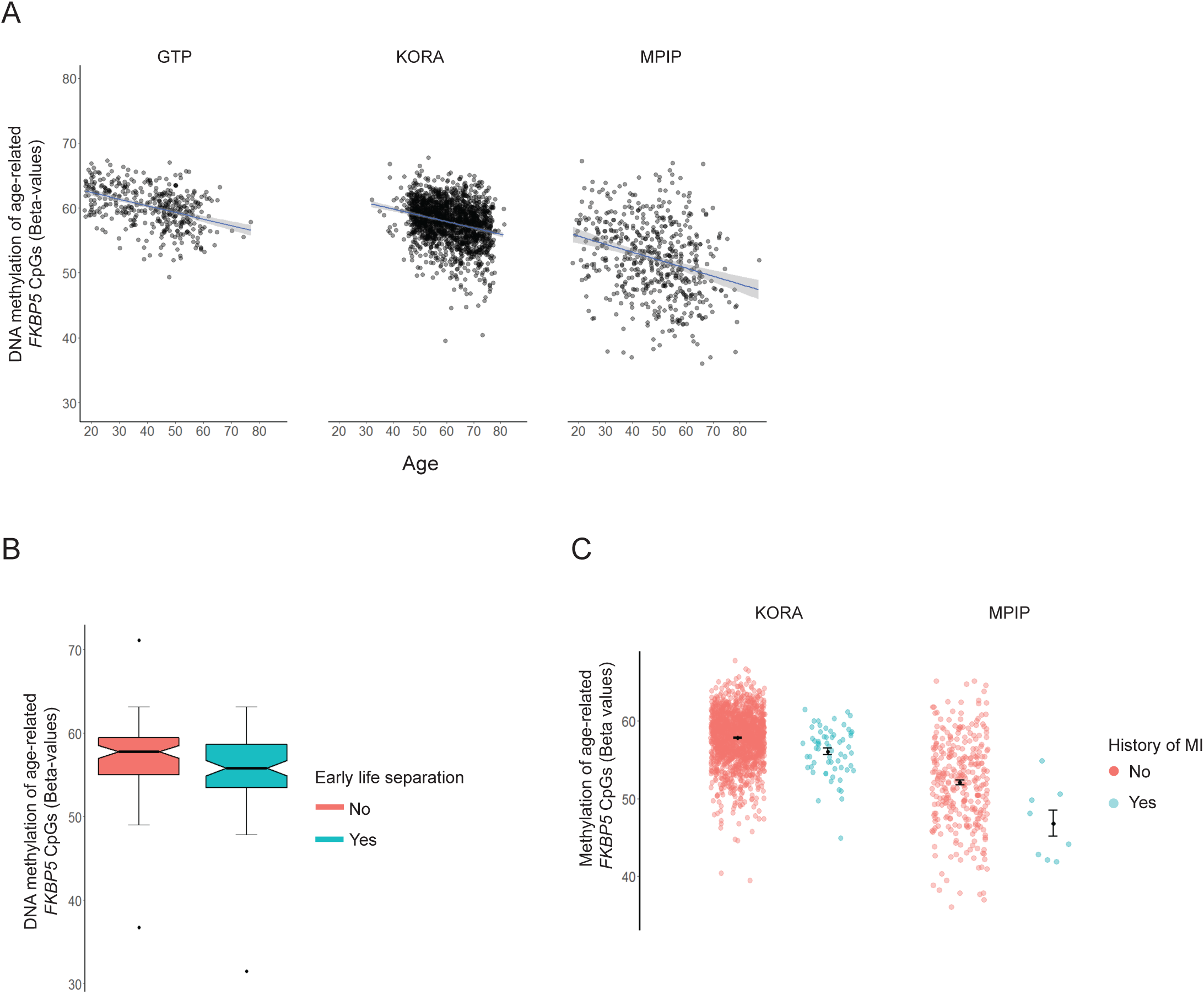
Complementary figure demonstrating how % *FKBP5* DNA methylation levels are associated with aging, early life separation, and history of myocardial infarction. The y axis in all panels depicts the average % DNA methylation (Beta-values) of the two age- and stress-related *FKBP5* CpGs (cg20813374 and cg00130530). Further details and statistics are provided in main Figures 1 and 6. GTP, Grady Trauma Project; HBCS, Helsinki Birth Cohort Study; KORA, Cooperative Health Research in the Region of Augsburg F4 community study; MI, myocardial infarction; MPIP, Max Planck Institute of Psychiatry depression case/control study.

**Supplementary Figure 2.**
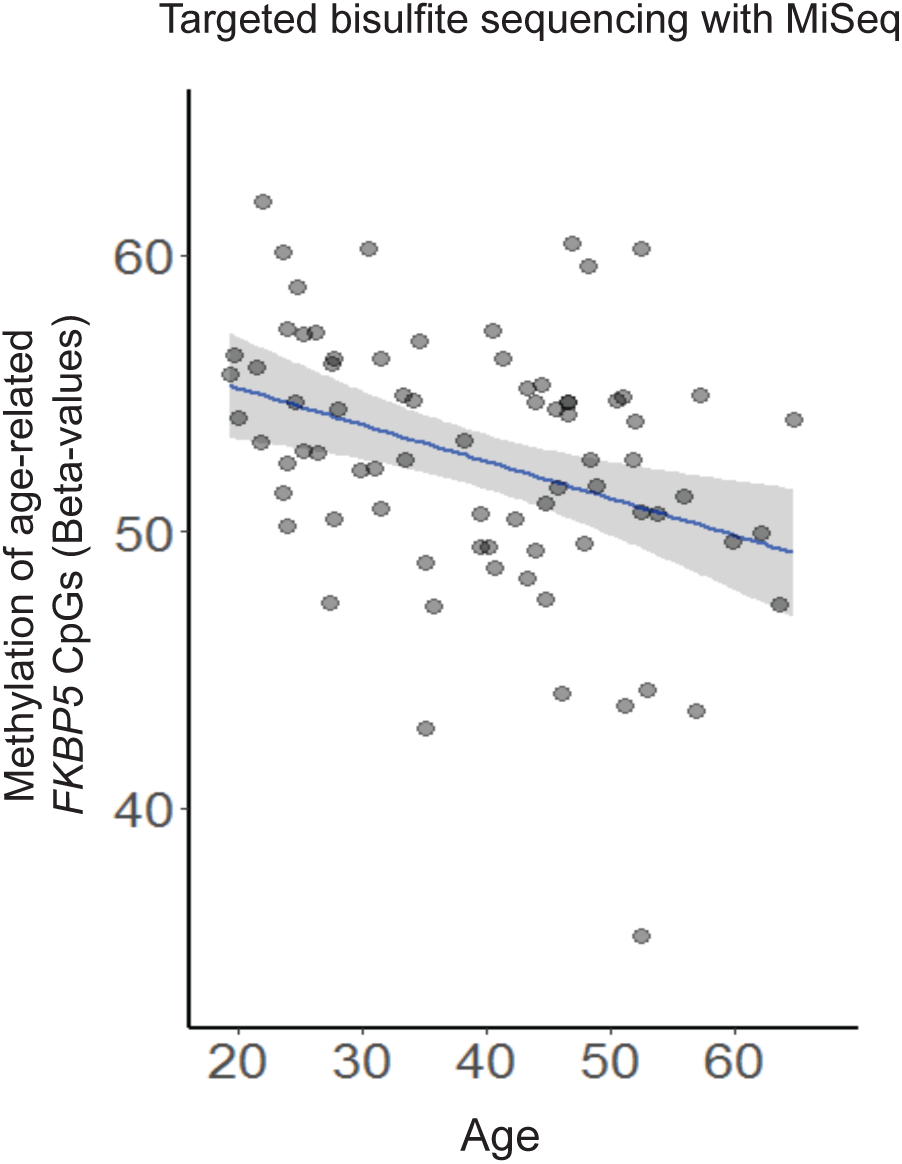
Validation of the 450K-identified age-regulated *FKBP5* CpGs (cg20813374 and cg00130530) using targeted bisulfite sequencing with the Illumina MiSeq in a sample of female subjects (n = 77, β_age_ = -0.0074, SE = 0.0031, p = 1.9 x 10^-2^). The y axis depicts the average % DNA methylation (Beta-values) of the two *FKBP5* CpGs. Reported statistics are after correcting for potential confounders (see Methods).

**Supplementary Figure 3.**
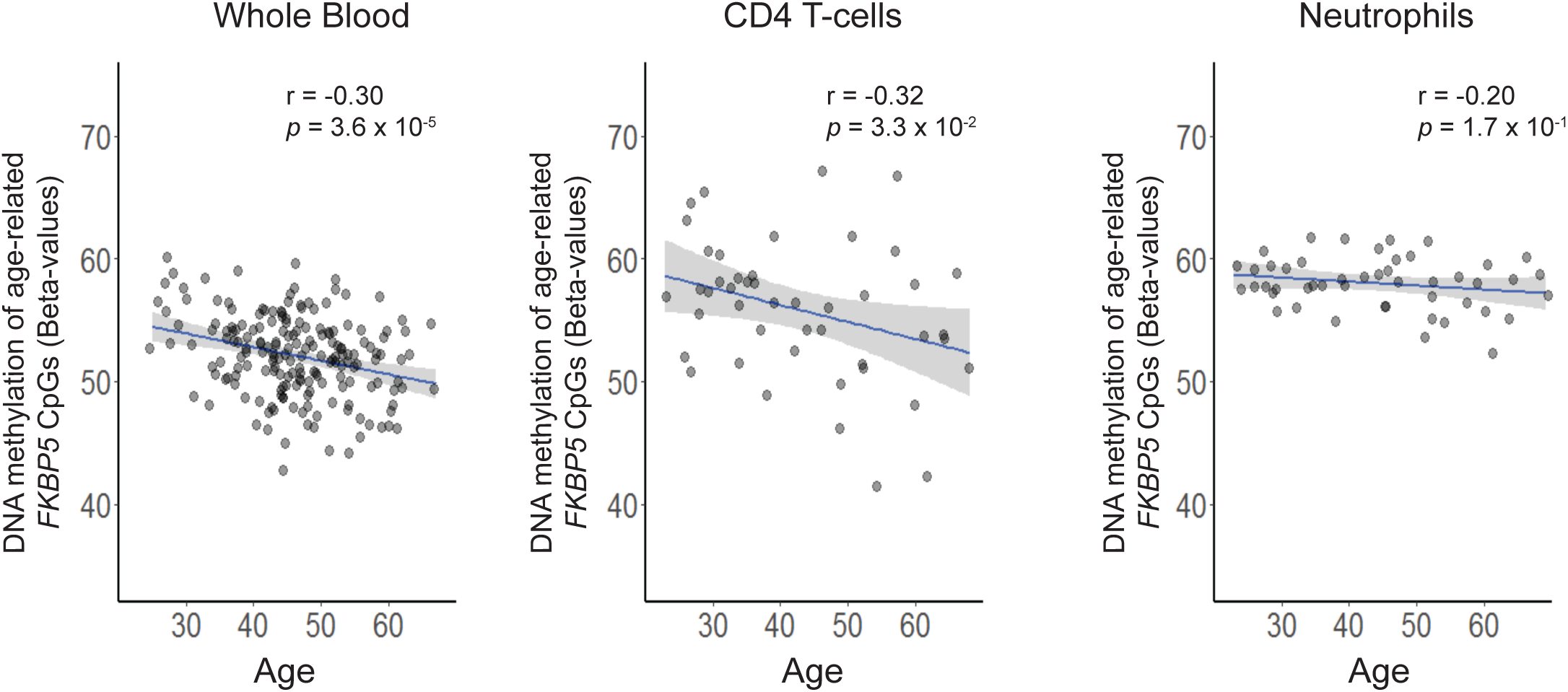
Pearson correlations between age and average methylation of the two age-regulated CpGs (cg20813374 and cg00130530) using publicly available DNA methylation data in whole blood (n = 184), as well as FACS-sorted CD4 T-cells (n = 46) and neutrophils (n = 48), from male subjects with a broad age range (source: GEO accession number, GSE67705, ref 43).

**Supplementary Figure 4.**
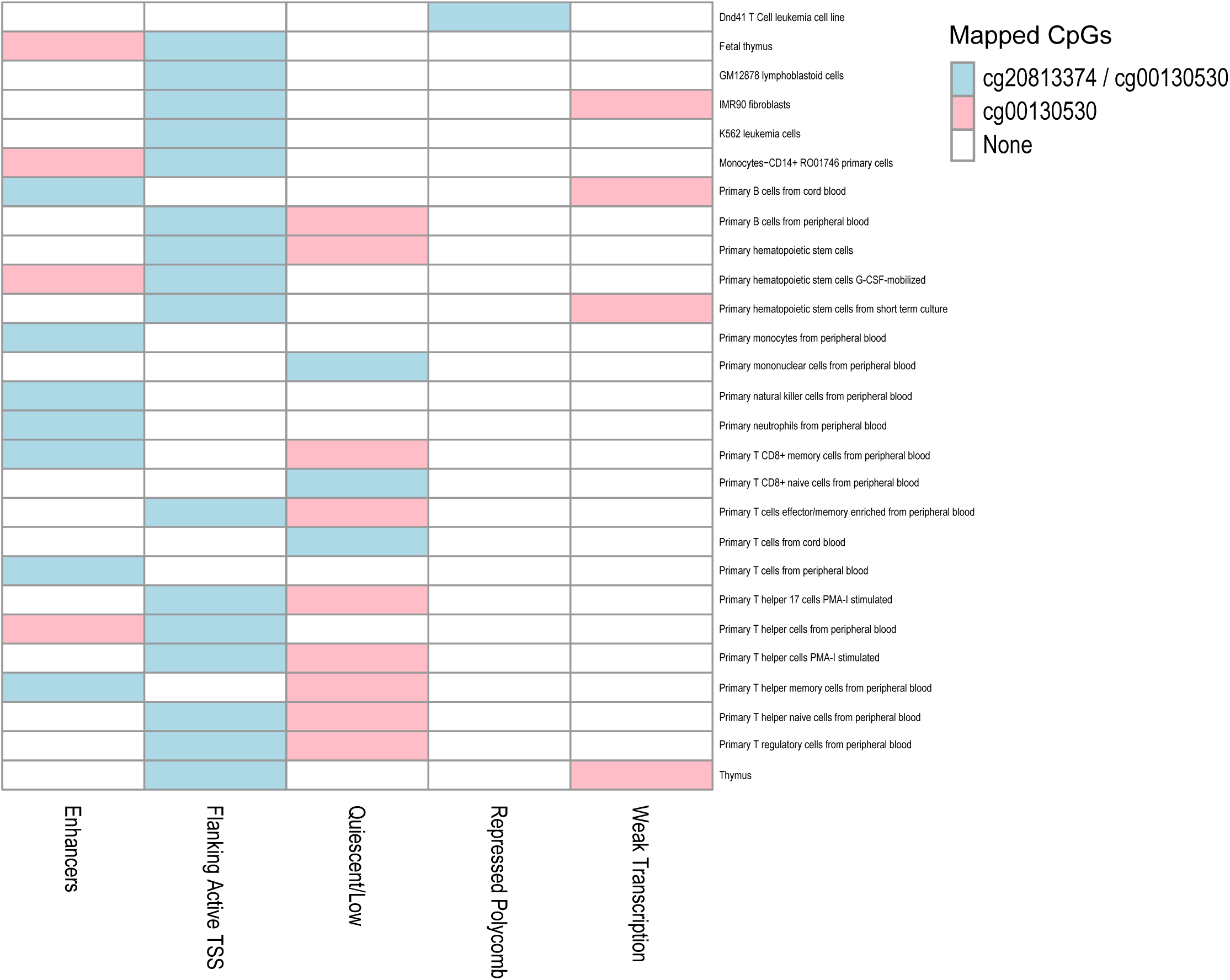
Visual depiction of integrative analysis of chromatin states using ChromHMM (ref 52) within immune cell types, as well as in the IMR-90 fibroblasts that were used as a model of human replicative senescence. Across these cell types, the age- and stress-related *FKBP5* CpGs are commonly mapped to an enhancer or flanking active TSS (further details in Supplementary Table 3).

**Supplementary Figure 5:**
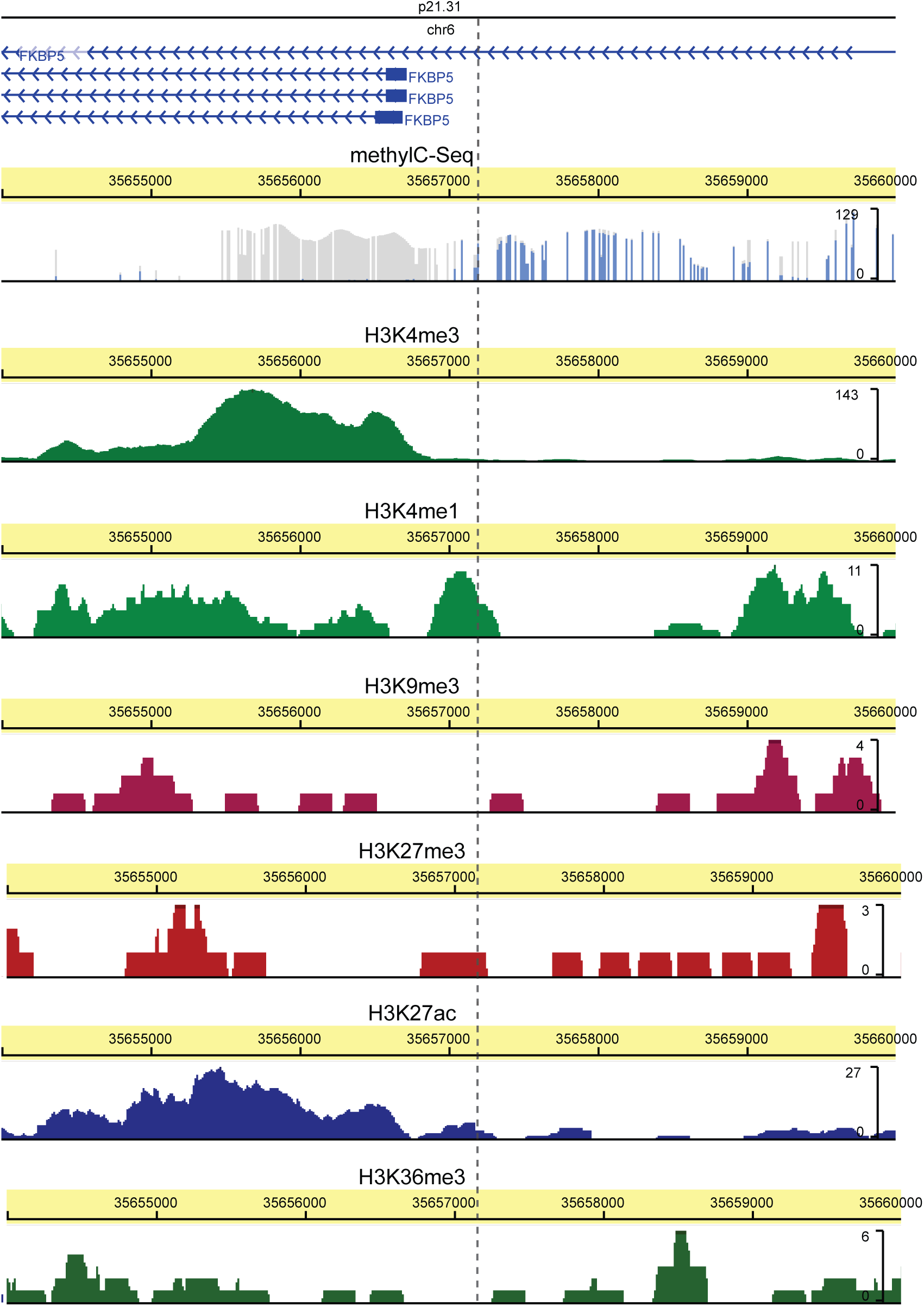
Functional annotation of the *FKBP5* locus at and around the age-related *FKBP5* CpGs, using the Roadmap Epigenome Browser and the “Mobilized CD34 Primary cells” track as a proxy for peripheral immune cells. The two CpGs (cg20813374 and cg00130530) are respectively located at positions 35657180 and 35657202 of chromosome 6 (exact location indicated by dotted line) and, as shown, exhibit intermediate methylation levels and colocalize with H3K4me1 and H3K27me3 signatures. This landscape is most consistent with a poised enhancer (ref 53).

**Supplementary Figure 6.**
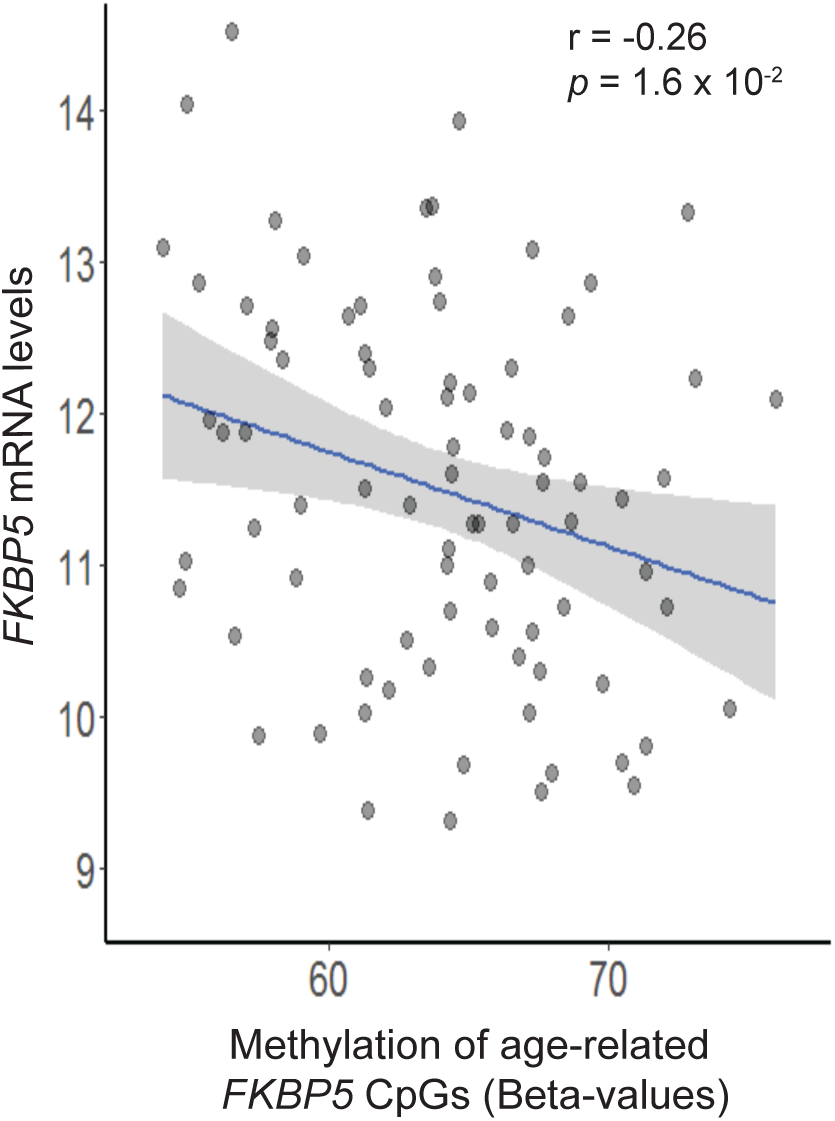
Methylation levels at the age- and stress-related *FKBP5* CpGs (cg20813374 and cg001305030) inversely correlate with *FKBP5* mRNA levels in breast tissue samples of control female subjects (n = 84). Publicly available data were analyzed from the Cancer Genome Atlas Wanderer (http://maplab.imppc.org/wanderer/; ref 54). The x axis depicts the average % DNA methylation (Beta-values) of the CpGs. The y axis depicts log2-transformed normalized RSEM RNAseq-measured *FKBP5* expression.

**Supplementary Figure 7.**
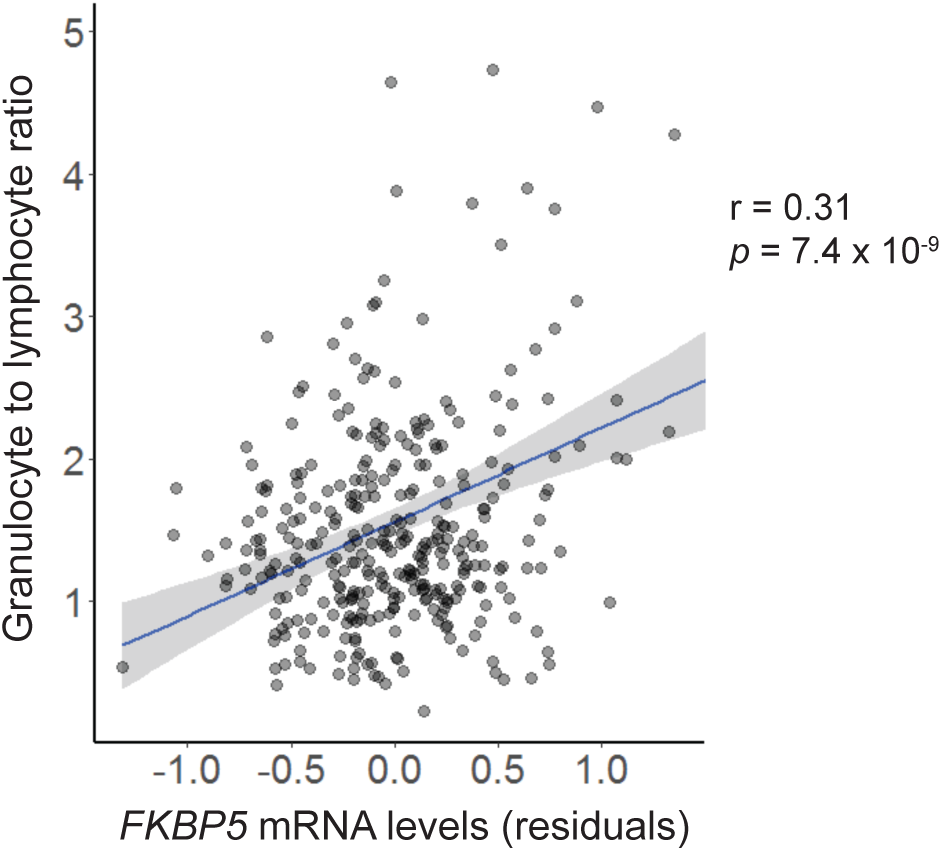
Association of *FKBP5* expression levels with the granulocyte to lymphocyte ratio (n = 330), an inflammation marker linked with heightened cardiovascular risk and mortality. Reported statistics and depicted residuals (on the x axis) are after correcting for covariates (see Methods).

**Supplementary Figure 8.**
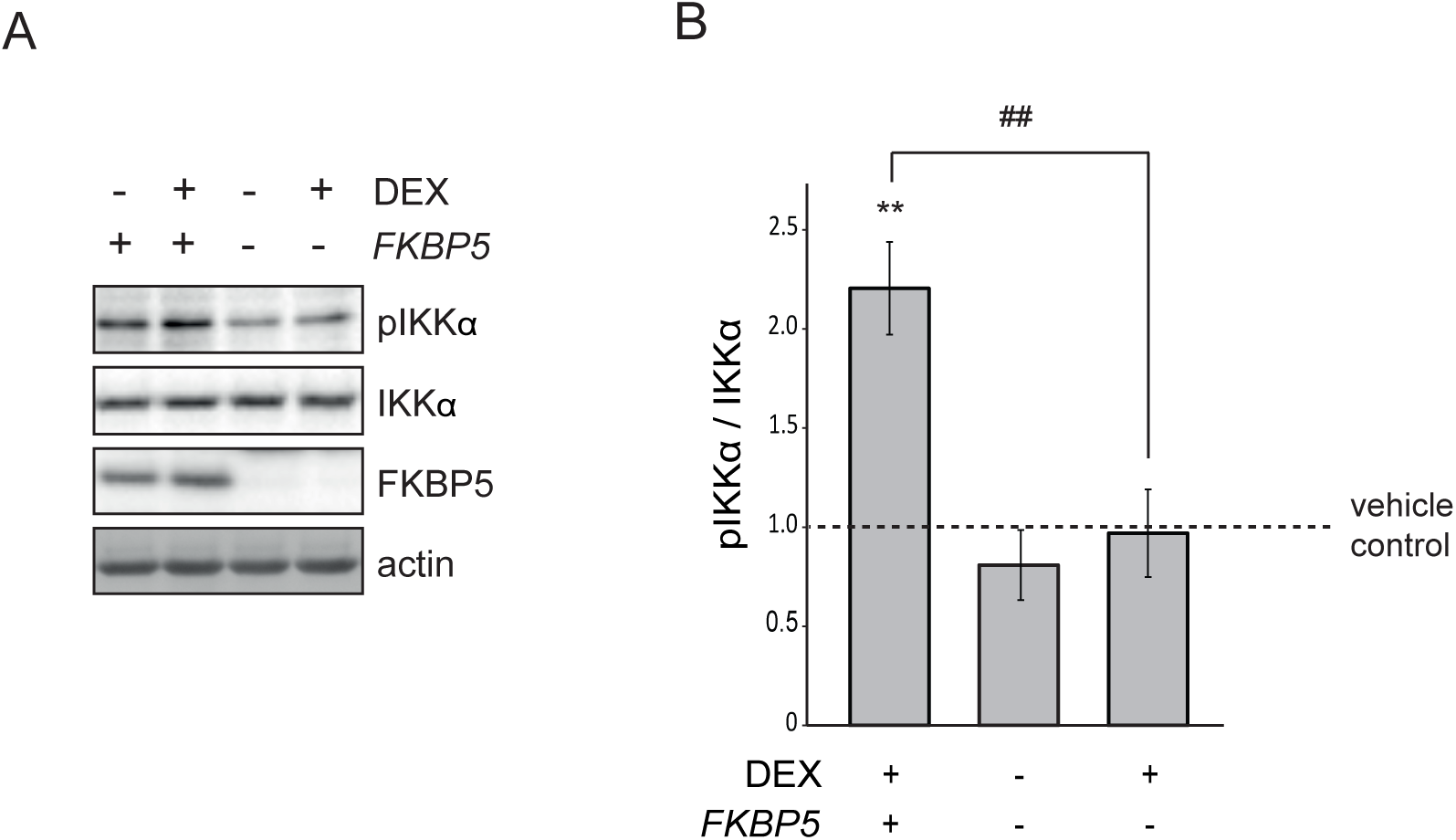
The effect of the stress hormone (glucocorticoid) receptor agonist dexamethasone (DEX) on functional phosphorylation of IKKα at serine 176 (pIKKα) is abolished in Jurkat cells lacking *FKBP5*. *FKBP5* knockout cells were generated using CRISPR/Cas9 technology. **(A)** Representative Western blots in lysates of Jurkat cells with or without the *FKBP5* gene treated for 24 hours with DEX (100nM), which robustly induces *FKBP5* expression, or vehicle (DMSO). (**B**) Western blots were quantified, and pIKKα levels were normalized to total IKKα levels and are shown as fold changes in comparison to wild-type (FKBP5 +) vehicle-treated cells. The treatment-genotype interaction was tested using two-way ANOVA (F_1,8_ = 8.1, interaction p = 2.2 x 10^-2^, n = 3 replicates per condition) and significant effects were followed with -2 Bonferroni-corrected pairwise comparisons. ** p < 10^-2^, statistically significant pairwise comparisons for vehicle *vs.* DEX. ## p < 10^-2^, statistically significant pairwise comparisons for cells with (wild-type) *vs.* cells without *FKBP5* (knockout).

**Supplementary Figure 9.**
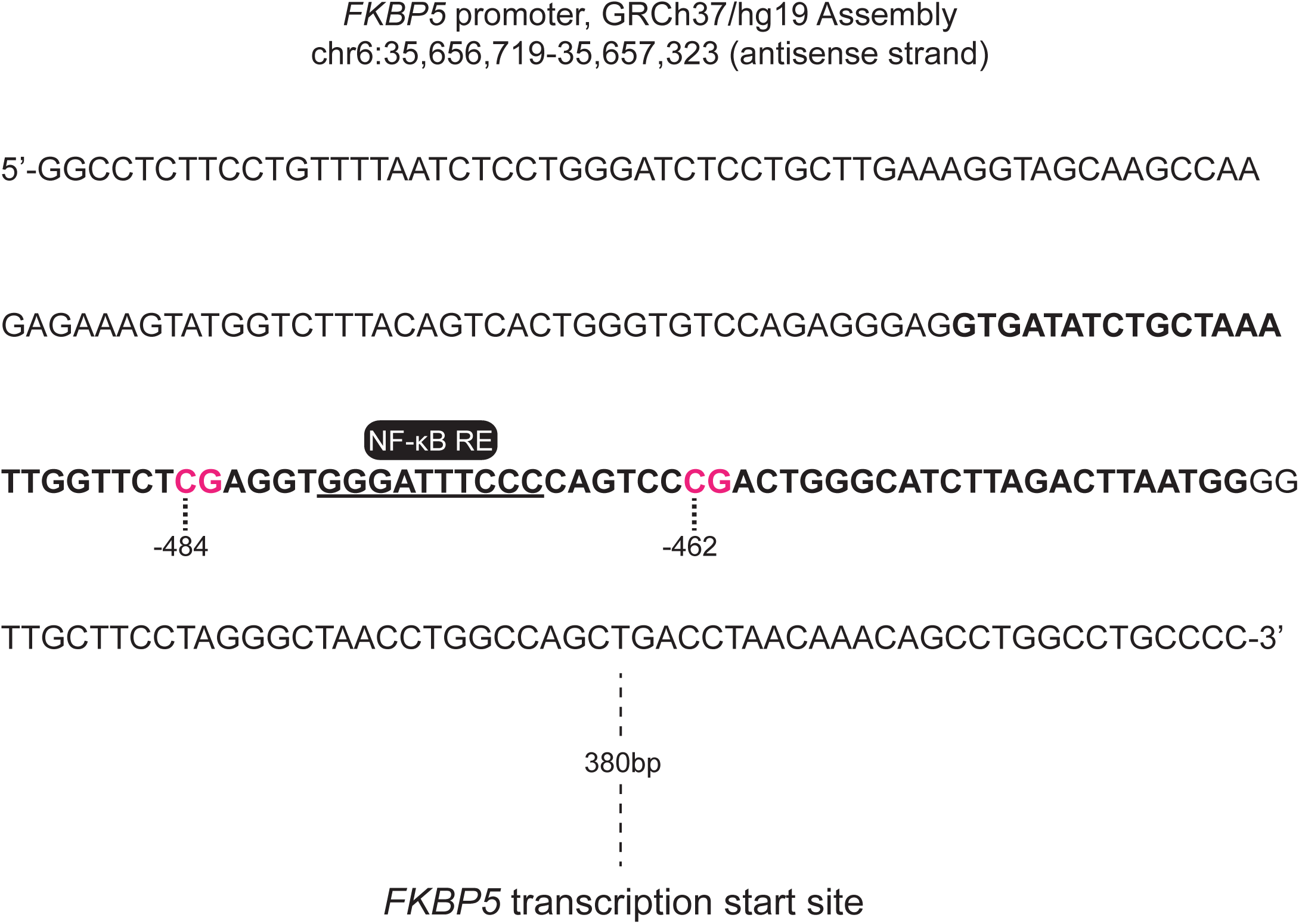
Annotation of the age- and stress-related *FKBP5* CpGs and the constructs used to characterize their function. The figure shows in detail the 5’-3’ sequence upstream of the *FKBP5* transcription start site of the DNA stretch (length 224 bp) that was inserted into the CpG-free luciferase reporter vector (ref 63). The black bold letters highlight the sequence of the biotinylated probe (length 70 bp) used for the biotinylated oligonucleotide-mediated chromatin immunoprecipitation. The pink letters highlight the age- and stress-related *FKBP5* CpGs. The underlined sequence and label highlight the NF-κB response element (NF-κB RE). As shown, both constructs include the CpGs and response element of interest, while they also completely lack other CpG sites to avoid non-specific CpG methylation effects.

